# mRNA poly(A)-tail length is a battleground for coronavirus–host competition

**DOI:** 10.1101/2025.10.09.680815

**Authors:** Arash Latifkar, Yevgen Levdansky, Amer Balabaki, Sherry S. Nyeo, Eugene Valkov, David P. Bartel

## Abstract

Most eukaryotic mRNAs contain a poly(A) tail, which in post-embryonic cells enhances their stability. Many cytoplasmic RNA viruses also harbor poly(A) tails on their genomic RNA and mRNAs. Here, we report that coronavirus infection causes cytoplasmic poly(A)-binding protein (PABPC) activity to become limiting, which preferentially destabilizes short-tailed host mRNAs, occurring before the action of virally encoded mRNA-decay factor nsp1. In this environment hostile to poly(A) tails, viral RNAs maintain a narrow tail-length distribution centering on 70–80 nucleotides across infection cycles. They do this through two mechanisms. First, viral tails are extended during RNA synthesis within double-membrane vesicles; second, viral tails are capped by a complex that includes PABPC1 and CSDE1 and slows tail shortening. Our findings suggest poly(A)-tail length is an arena of host– virus conflict, in which preserving tail lengths of viral mRNAs promotes their cytoplasmic dominance.

**Highlights:**

- PABPC1 becomes limiting during coronavirus infection
- Limiting PABPC1 promotes decay of short-tailed host mRNAs—independently of nsp1
- The tail lengths of coronaviral mRNAs are extended during their synthesis in DMVs
- Viral tails are capped by PABPC1 and CSDE1, which protects against deadenylation

## Introduction

The poly(A) tail, a stretch of adenosines at the 3′ end of an mRNA, is a hallmark feature of nearly all eukaryotic mRNAs. In these eukaryotes, the poly(A) tail is generated co-transcriptionally in the nucleus by nuclear poly(A) polymerase enzymes, which synthesize poly(A) tail with average lengths of ∼200 nucleotides (nt) in mammals^1^ and ∼70 nt in yeast^2^.

Many eukaryotic viruses also carry poly(A) tails on their mRNAs, and in some RNA viruses, also on their genomes. DNA viruses that replicate and transcribe their genome in the nucleus can access host nuclear polyadenylation machinery to generate poly(A) tails. In contrast, for some that are RNA viruses, the poly(A) tail of viral mRNAs is templated in (–) strand RNA with a stretch of poly(U)s. Such templated regeneration of the poly(A) tail poses a unique challenge for these cytoplasmic RNA viruses that have polyadenylated mRNA genomes. They rely on the length of the poly(U) template generated from the poly(A) tail of a genomic (+) strand transcript to generate the poly(A)-tail length in the progeny mRNAs. In this scenario, cytoplasmic deadenylation by the PAN2– PAN3 and CCR4–NOT complexes^3^ is expected to cause progressive shortening of poly(A) tails over multiple serial infections. For example, even under optimal conditions where viral mRNAs experience one of the slowest measured deadenylation rates, such as the 0.15 nt/min rate of TOP mRNAs^4^, and are deadenylated for only one-hour post-cytoplasmic entry, viral mRNAs would lose more than half of their initial poly(A)-tail length during five serial infections (**Figures 1A and S1**). We refer to this challenge as the “tailomere problem,” drawing an analogy to the telomere problem in DNA replication.

**Figure 1.**
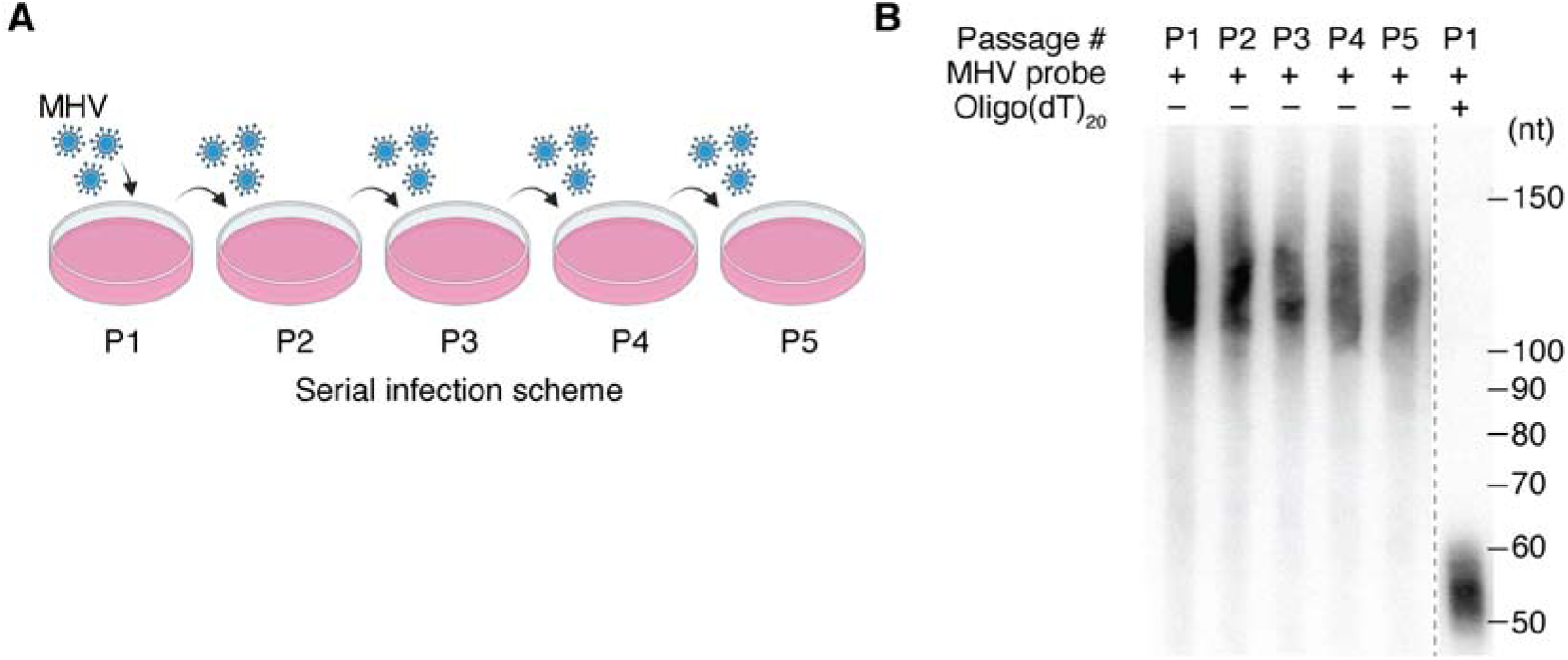
Poly(A)-tail length of viral mRNAs remains unchanged during serial infection passages. (A) Outline of serial infection with mouse hepatitis virus (MHV). For each round of infection, the progeny virus is used to infect a subsequent set of uninfected cells. (B) Unchanged poly(A)-tail length of viral mRNAs during serial infection. Shown is an RNase H northern blot that measures poly(A)-tail lengths of viral mRNAs in virions for each passage of a serial infection in NIH-3T3 cells.

Viruses have been proposed to employ several mechanisms, including ‘stuttering’ of the viral RNA polymerase on poly(U)s and nontemplated nucleotidyl transferase activity of the viral RNA polymerase or host polyadenylation machinery, to counteract the tailomere problem^5–9^. Despite these existing models, how cytoplasmic deadenylation influences the poly(A) tail of viral mRNAs is poorly understood, as are the determinants of poly(A)-tail length for viral mRNAs.

Maintaining the poly(A)-tail length of an RNA-based viral genome is presumably essential to viral fitness for many reasons. Poly(A)-tail length plays a critical role in regulating mRNA translation and stability. In oocytes and early embryos, poly(A)-tail length is tightly coupled to translation efficiency, whereas in post-embryonic cells, tail length has little effect on translation and instead influences mRNA stability^4,10,11^. For viruses, early studies in poliovirus demonstrated that removing the poly(A) tail from the viral genome drastically attenuates infectivity^12^. Further research has revealed that the poly(A) tail can act as a crucial cis-acting signal for viral RNA replication and that a poly(A) tail of at least 12 nt is necessary for efficient negative-strand RNA synthesis and infectivity^13,14^. Similarly, in coronaviruses, the poly(A) tail is essential for RNA replication, with its length directly impacting the efficiency of replication. Defective genomes with a missing or shortened poly(A) tail have substantially reduced or delayed replication, although some viruses can restore the tail-length over time^15,16^. Here, we investigated the maintenance and extension of poly(A) tails in betacoronavirus mRNAs, focusing on Mouse Hepatitis Virus (MHV).

## Results

### Poly(A)-tail length of viral mRNAs remains unchanged during serial infection passages

Like the SARS-CoV-2 virus, mouse hepatitis virus is a betacoronavirus with a positive-sense, single-stranded, polyadenylated RNA genome. Prior studies show that the poly(A)-tail length of viral mRNAs is an important determinant of MHV infectivity^15,16^. We found that the tail length of MHV mRNAs was unchanged during serial infection that spanned five passages (**Figure 1B**), prompting us to further investigate the dynamics of poly(A) tails in both host and viral mRNAs during infection. Previous studies examining poly(A)-tail dynamics during coronavirus infection were conducted at timepoints that encompass multiple rounds of infection ^17–19^. To gain a better understanding of early regulatory events, we focused our analysis on timepoints spanning a single round of infection.

### Poly(A)-tail extension of viral mRNAs occurs in double membrane vesicles

To better understand poly(A)-tail metabolism during viral infection, we infected mouse L2 cells with MHV and conducted Poly(A)-tail Length sequencing (PAL-seq)^10^ on RNA extracted from the cytoplasmic fraction of the infected cells (**Figure S2A**). Fractionation efficiency was confirmed by marker distribution: PABPC1 localized to cytoplasm, Histone H3 to nucleus, and Xbp1 splicing products^20^ segregated as expected (**Figure S2B–C**).

Viral mRNA abundance had the anticipated dynamics, with cytoplasmic levels of viral mRNAs dramatically increasing as the infection progressed (**Figure 2A**). Interestingly, the median poly(A)-tail length of viral mRNAs increased after their initial appearance in the cytoplasm (76 nt), reaching a peak median length of 85 nt at 6 hours post-infection (hpi) (**Figure 2B**, **top**). Importantly, the poly(A)-tail length distribution of zebrafish mRNAs, spiked into each RNA sample as a control, remained unchanged, which confirmed the reproducibility of our measurements (**Figure 2B**, **bottom**). The observation that tail lengths increased over the first 6 h aligned with previous reports on bovine coronavirus (BCoV)^21^ and avian infectious bronchitis virus (IBV)^22^, which described an extension of viral poly(A)-tail lengths during the early hours following viral entry. However, the mechanism behind this poly(A)-tail extension has remained elusive.

**Figure 2.**
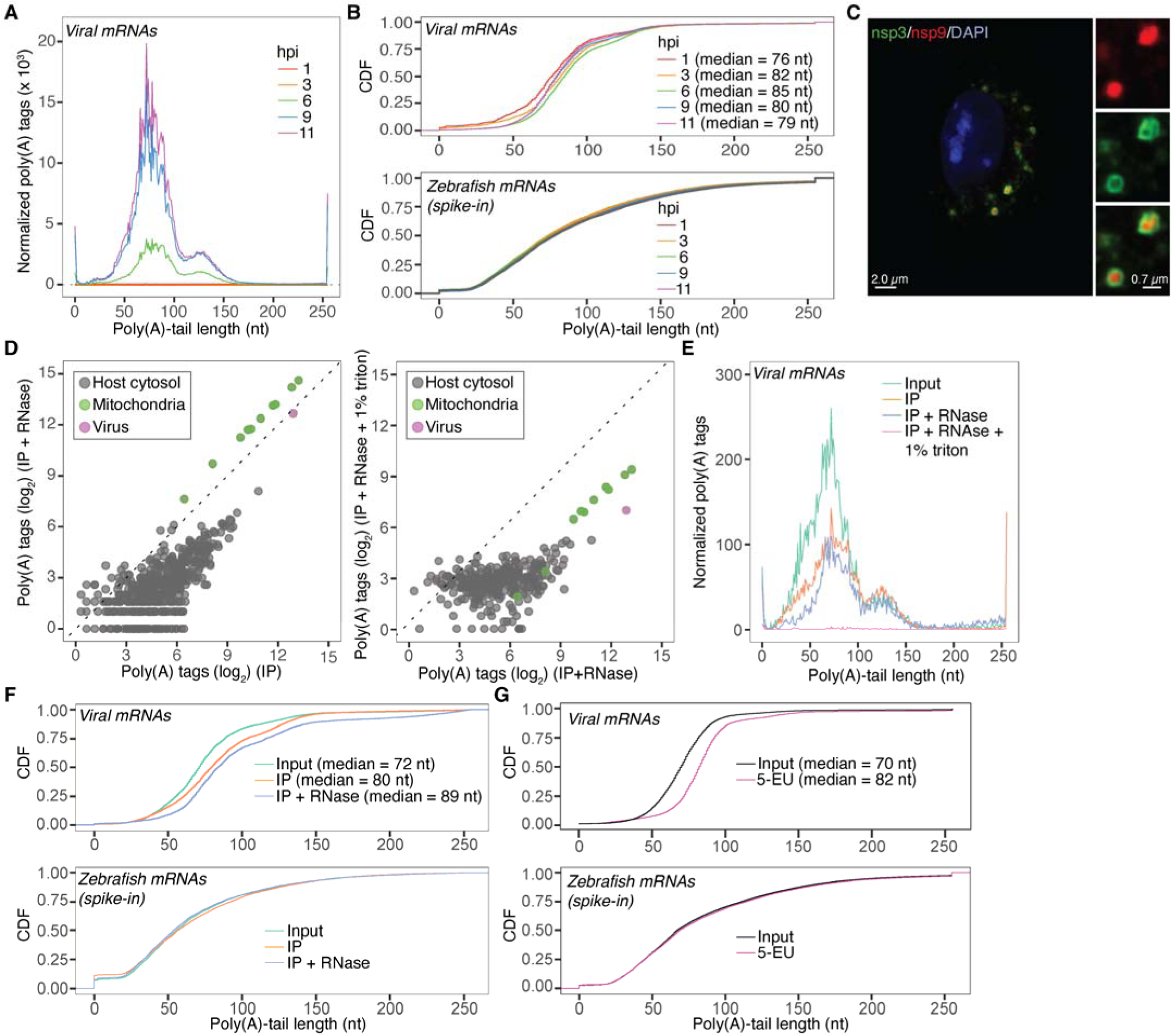
Poly(A)-tail extension of viral mRNAs occurs in double membrane vesicles. (A) Progressive increase in viral mRNA levels during infection. Plotted are normalized tail-length abundances of viral mRNAs in mock and MHV-infected cells. (B) Increase in poly(A)-tail lengths of viral mRNAs during infection. Plotted are Cumulative distribution functions (CDFs) of the poly(A)-tail lengths of viral mRNAs (top), and of zebrafish mRNAs (bottom) that were spiked in as internal controls. (C) Immunofluorescent image of a cell infected with MHV-Δ2-GFP3 strain^30^. The DMVs are marked by GFP-tagged nsp3 and the viral RdRps are marked by nsp9. (D) Membrane-protected mRNAs enriched in an IP of DMVs. *Left*, Effect of RNase1 on mRNAs that co-purified with DMVs. Shown are the abundances of mRNAs in GFP immunoprecipitates treated with RNase I, plotted in relation to their abundance in the same precipitates that were not treated. *Right*, Effect of detergent in exposing mRNAs that co-purified with and were protected by DMVs. Plotted are the abundances of mRNAs in GFP immunoprecipitates treated with triton prior to RNase I treatment compared to the same precipitates that were treated only with RNAse I. (E) Tail-length distributions of viral mRNAs co-purifying with DMVs. Plotted are normalized tail-length abundances of 5 (E) Tail-length distributions of viral mRNAs co-purifying with DMVs. Plotted are normalized tail-length abundances of viral mRNAs in cytoplasmic input and DMV IPs. (F) Tail-length distributions of viral mRNAs co-purifying with DMVs. Plotted are CDFs of the poly(A)-tail lengths of viral mRNAs (top) and of zebrafish mRNAs (bottom) that were spiked in as internal controls. (G) Tail-length distributions of bulk and nascent viral mRNAs. Plotted are CDFs of poly(A)-tail lengths of total and nascent viral mRNAs (top, input and 5-EU, respectively) and of zebrafish mRNAs (bottom) that were spiked in as internal controls.

Viral replication organelles are double-membrane vesicles (DMVs) formed through remodeling of the endoplasmic reticulum (ER). These structures are generated by nonstructural proteins translated from the viral genomic mRNA. They are associated with the viral RdRP and serve as sites of viral RNA synthesis^23^. Noting the temporal overlap between the extension of poly(A) tails on viral mRNAs and the formation of these DMVs^24–26^, we examined whether poly(A)-tail extension is linked to viral mRNA synthesis within these compartments. Accordingly, we used organelle immunoprecipitation^27–29^ (IP) to isolate DMVs during infection, taking advantage of EGFP-tagged nsp3^30^, a viral protein integral to the pore structure on DMVs^31,32^ (**Figure S2D**). From these DMVs immunoprecipitated during infection, RNA was isolated and tail lengths profiled using PAL-seq.

Several tests confirmed the quality of our DMV IP. Immunofluorescent microscopy confirmed that nsp3-GFP co-localized with the nsp9 subunit of the viral RdRp (**Figure 2C**) and that our pulldowns enriched for viral RdRp (mature nsp9) while excluding cytosolic (PABPC1) and non-ER organellar markers (**Figure S2E**). Consistent with DMVs originating from the ER, our IPs were enriched for the ER marker Calreticulin (CALR). To ensure focus on mRNAs within DMVs, we treated the immunoprecipitates with RNAse I, which reduced the abundance of most cytoplasmic mRNAs, causing relative enrichment of viral and mitochondrial mRNAs, the latter also residing in membrane-protected environments (**Figure 2D, left**). Detergent treatment (1% Triton) caused degradation of viral and mitochondrial mRNAs together with cytoplasmic RNA, as expected if the viral and mitochondrial RNAs had been protected by their respective membranes (**Figure 2D, right**).

Tail-length profiling revealed that membrane-protected viral mRNAs had significantly longer tails compared to bulk cytoplasmic viral mRNAs (**Figure 2E** and **Figure 2F, top**), whereas zebrafish mRNA spiked in as an internal control for measurement reproducibility remained unaffected (**Figure 2F, bottom**). The observation that DMVs contained viral mRNAs that had longer tail lengths raised the question of whether poly(A)-tail extension occurs during viral mRNA synthesis. Leveraging a recent report that 5-ethynyl uridine (5-EU) labels nascent viral mRNAs within DMVs^33^, we set out to answer this question. Infected cells were pulsed with 5-EU, followed by RNA isolation, biotin click chemistry, streptavidin capture, and poly(A)-tail profiling (**Figure S2D**). As expected^34^, nascent mitochondrial mRNAs had shorter poly(A)-tails compared to the bulk population (**Figure S2F**), which validated this approach. In contrast, nascent viral mRNAs had substantially longer poly(A)-tails (**Figure 2G, top**), whereas zebrafish mRNA spike-ins remained unchanged (**Figure 2G, bottom**).

Together, our findings strongly support the conclusion that viral mRNA poly(A)-tail extension occurs during synthesis within the membrane-protected environment of DMVs. These observations were consistent with reports showing that MHV defective interfering (DI) RNAs—lacking or bearing shortened poly(A) tails—can restore their tails through a replication-coupled repair mechanism^15,16^.

### A host RBP forms a complex with viral poly(A) tail and PABPC1

The analysis of poly(A)-tail lengths in viral mRNAs uncovered a striking feature compared to cytoplasmic host mRNAs. Specifically, the distribution of poly(A)-tail lengths in viral mRNAs was much narrower (**Figure 3A**). When comparing the standard deviation of poly(A)-tail lengths for individual genes, viral mRNAs resembled mitochondrial mRNAs more than they resembled cellular mRNAs (**Figure 3B**). A similar result was observed for SARS-CoV-2 mRNAs (**Figure 3C**), consistent with previous reports of poly(A)-tail lengths in SARS-CoV-2-infected cells^19^. Because mitochondrial mRNAs are membrane protected and are not subject to deadenylation by the cytoplasmic deadenylation machinery, CCR4–NOT, we wondered whether viral mRNAs might also be protected from deadenylation. Because the narrow distribution was shared between SARS-CoV-2 and MHV, we leveraged available datasets that report RNA-binding proteins (RBPs) that directly interact with the viral mRNAs^35,36^, with the idea that some RBPs might confer resistance to deadenylation. Consistent with prior reports^16^, cytoplasmic poly(A)-binding proteins (PABPC1 and PABPC4), were identified as binding to viral mRNAs (**Figure S3A**). Because PABPCs can act to stimulate deadenylation^37,38^, we searched for other RBPs that might form a complex with PABPCs to slow deadenylation. To this end, we performed an *in silico* screen using AlphaFold3 (AF3)^39^, modeling potential interactions between PABPC1 and host or viral RBPs. The screen was conducted using 96 RBPs that interact with viral mRNAs (virus-associated RBPs) and either full-length PABPC1 or its individual domains: RRM1–2, RRM3–4, and the C-terminal PABC domain. To assess the predicted interaction propensity between PABPC1 and the virus-associated RBPs, we applied the PEAKscore metric^40^, which quantifies the Predicted Alignment Error (PAE) exclusively between the bait and the candidate protein, while excluding internal PAE within either protein. This screen recovered several previously reported interactions, including those between PABPC1 and either LARP1^41^, ATXN2^42,43^, or ATXN2L^44^, each of which binds the PABC domain of PABPC1 through a PAM2 motif^45,46^ (**Figure 3D**). In addition, the cold-shock-domain-containing protein E1 (CSDE1) ranked among the top three interactors with full-length PABPC1 and had the highest PEAKscore when queried with the RRM1–2 domain (**Figure 3D**). Specifically, the AF3 model predicts that the truncated cold-shock domain, CSD2, of CSDE1 directly interacts with RRM1 of PABPC1 (**Figures 3E and 3F**). As CSDE1 has been identified as a proviral protein in several studies^35,36,47^ and CRISPR screens^48,49^ (**Figure S3B**), we set out to better understand this predicted interaction.

**Figure 3.**
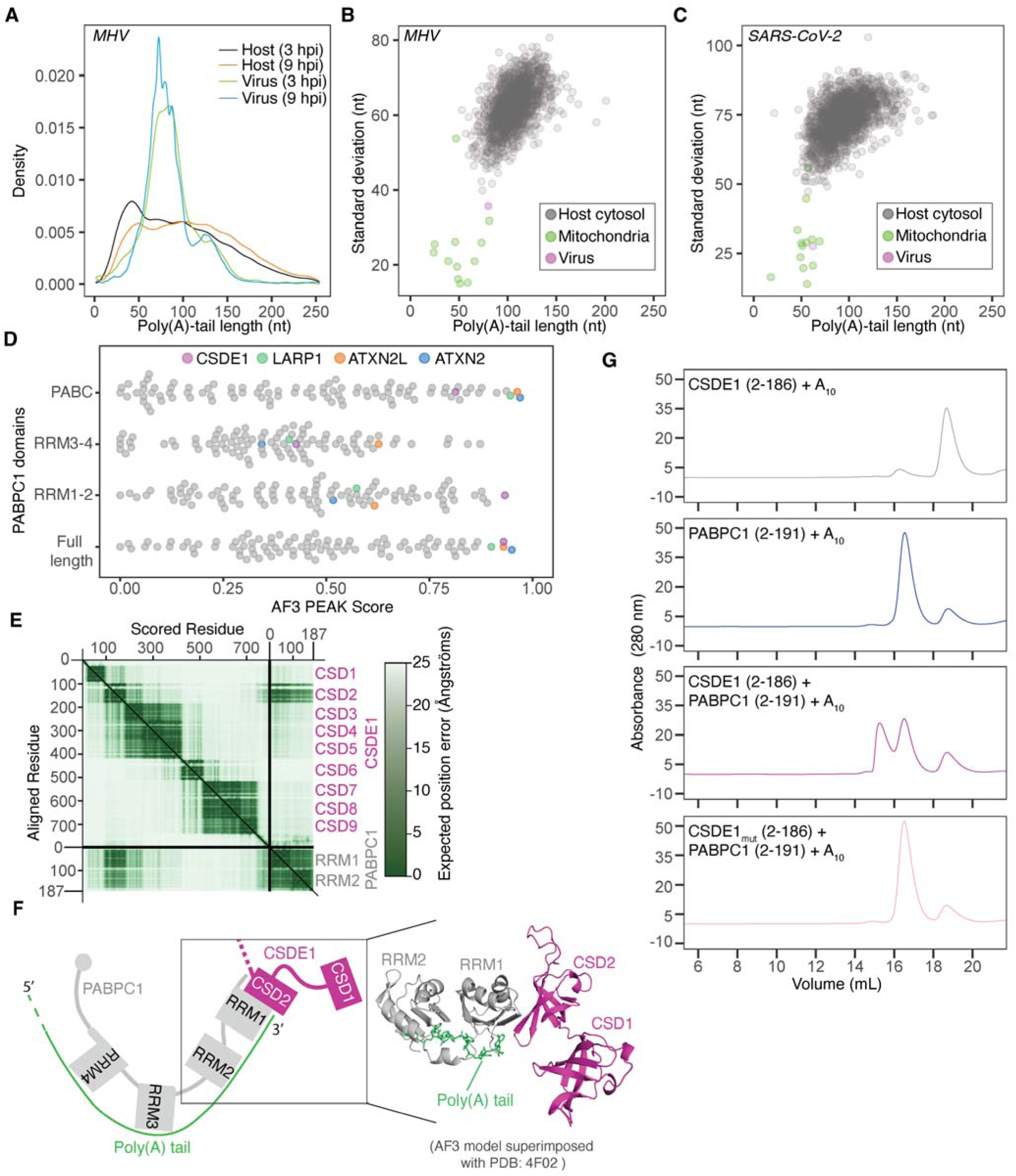
A host RBP forms a complex with viral poly(A) tail and PABPC1. (A) Narrow tail-length distribution of viral mRNAs compared to host mRNAs. Plotted are tail-length distributions of viral mRNAs (at 3 and 9 hpi) and host mRNAs (at 3 and 9 hpi) upon infection. (B) Narrow tail-length distribution of MHV mRNAs compared to host mRNAs. Shown for cytoplasmic mRNAs of virus or each host gene passing the expression cutoff are standard deviations for tail lengths in MHV-infected cells (at 9 hpi) plotted as a function of median tail length. (C) Narrow distribution of SARS-CoV-2 mRNAs compared to host mRNAs at 6 hpi; otherwise, as in (B). (D) AF3 prediction of multimers between virus-associated RBPs and PABPC1. Plotted are PEAK scores for an AF3 screen of multimers of virus-associated mRNA-binding proteins and either full-length PABPC1 or its individual domains. (E) The PAE plot generated from the AF3 model of the multimer between CSDE1 and RRM1-RRM2 domain of PABPC1. (F) Schematic and the 3D model of AF3-predicted complex between CSDE1 and RRM1-RRM2 domain of PABPC1. The poly(A) tail in the 3D model (right) is inserted from the superimposed structures of AF3 model and PDB: 4F02. (G) Biochemical support for the AF3 model of the complex between CSDE1 and RRM1-RRM2 domain of PABPC1. Shown are size-exclusion chromatograms of the minimal components of the complex: Residues 2-186 of CSDE1 comprising CSD1 and CSD2, residues 2-191 of PABPC1 comprising RRM1 and RRM2 of PABPC1, and oligo(A)_10_. The CSDE1_mut_ harbors mutations (V136R and Y138R) predicted to disrupt the interface between CSDE1 and the RRM1-RRM2 domain of PABPC1.

To test whether purified CSDE1 can form a complex with PABPC1–poly(A), we used size-exclusion chromatography, which showed that CSDE1, PABPC1, and an oligo(A)_30_ can form a ternary complex in vitro (**Figure S3C**). This result is consistent with prior studies describing CSDE1 as a binding partner for PABPC1^51,52^; however, in contrast to one of those studies^52^, we did not observe a complex between PABPC1 and CSDE1 in the absence of oligo(A)_30_ (**Figure S3D**). Moreover, the interaction mode predicted by AF3 differs from that of a prior report that mapped the CSDE1– PABPC1 interaction to the end of RRM2 and the entirety of RRM3 (amino acids 166–289) of PABPC1^52^. Notably, when we expanded the query to all RRM domains in the human proteome, the RRM1 of PABPC proteins were the top predicted interactors of CSD2 of CSDE1 (**Figure S3F**). To validate the AF3 prediction, we purified the minimal predicted interacting domains—CSD1 and CSD2 (residues 2–186) of CSDE1 and RRM1 and RRM2 (residues 2–191) of PABPC1—and tested their binding. These minimal domains, together with an oligo(A)_10_ formed a ternary complex, and this complex was disrupted when the residues at the AF3-predicted interface, namely V136 and Y138 of CSD2, were mutated (**Figure 3G**). Moreover, as measured by mass photometry, the ternary complex between PABPC1, CSDE1, and oligo(A)_30_ had a molecular weight of ∼150 kDa, which was consistent with the predicted 1:1:1 stoichiometry of the three components (**Figure S3E**).

### The PABPC1-CSDE1 complex protects viral mRNA from cytoplasmic deadenylation

PABPC1 is reported to stimulate deadenylation by the CCR4–NOT complex^37,38^. The AF3 model of PABPC1–CSDE1 interaction suggested that CSDE1 can limit the accessibility of the 3’ end of the poly(A)-tail to deadenylases by binding to the lateral face of RRM1 on PABPC1. To test this hypothesis, we examined the effect of CSDE1 on the in vitro deadenylation of an RNA substrate with an 80-nt poly(A) tail, matching the tail length of nascent viral mRNAs isolated by 5-EU labeling. We observed that addition of either the minimal (**Figure 4A, top**) or full-length CSDE1 (**Figure 4B, top**) construct slowed deadenylation compared to PABPC1 alone. Interestingly, although mutations that disrupted the interaction between PABPC1 and minimal CSDE1 completely abrogated the inhibition of deadenylation (**Figure 4A, bottom**), the full-length mutant CSDE1 retained the ability to slow deadenylation, achieving levels of inhibition intermediate between PABPC1 alone and the combination of PABPC1 with wild-type CSDE1 (**Figure 4B, bottom**).

**Figure 4.**
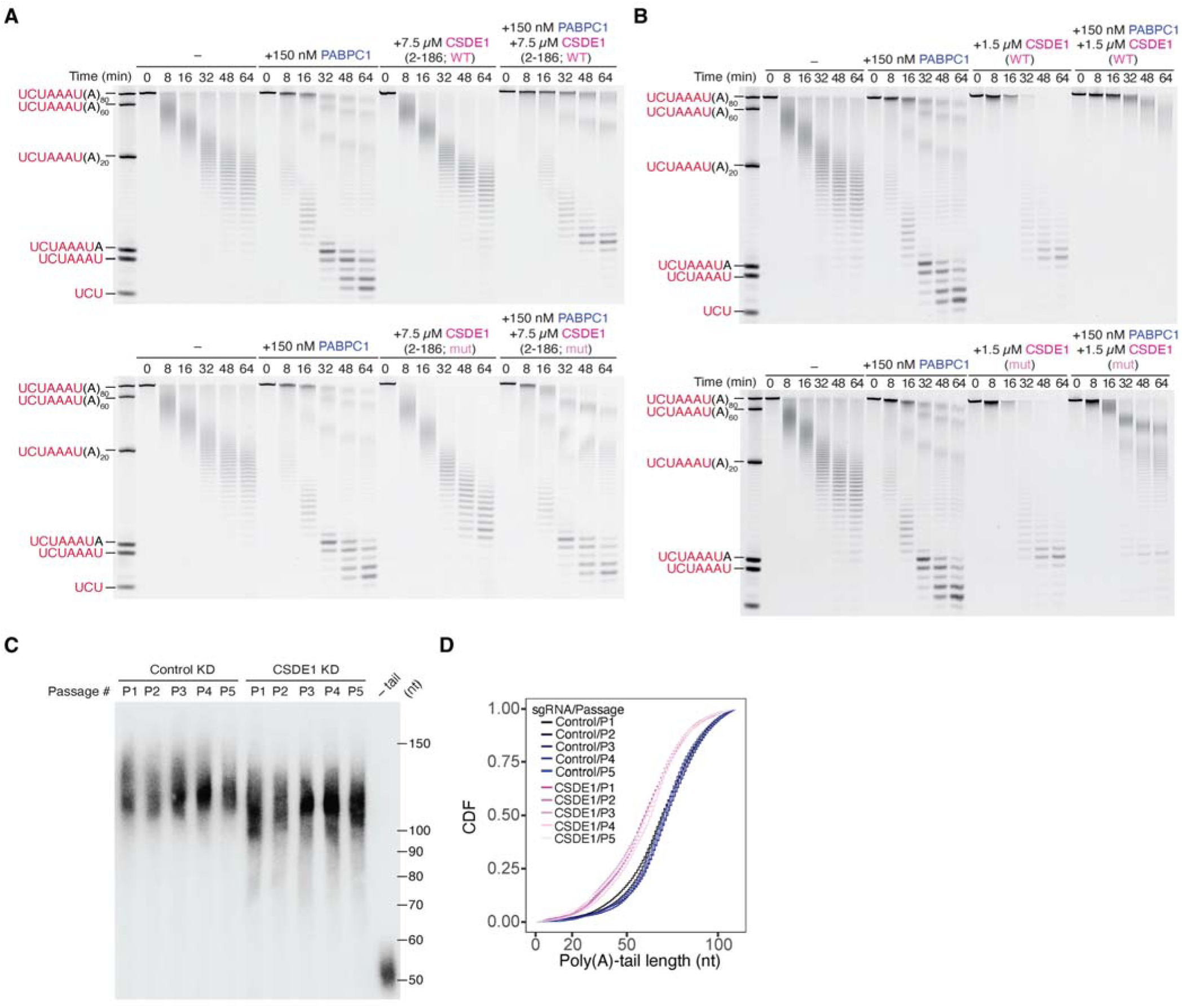
The PABPC1–CSDE1 complex protects viral mRNA from cytoplasmic deadenylation. (A) Repression of deadenylation by CSD1–2. Shown are time courses of in vitro deadenylation of a UCUAAAU(A)DD substrate by the full CCR4–NOT complex. Reactions were performed in the presence of either no added protein (–), PABPC1 only, CSDE1 residues 2–186 (containing CSD1 and CSD2) only in either wild-type (WT, top panel) or mutant (mut, bottom panel) form, or PABPC1 together with CSDE1 residues 2–186 in either wild-type (WT, top panel) or mutant (mut, bottom panel) form. (B) Repression of deadenylation by full-length CSDE1. Shown are time courses of in vitro deadenylation of a UCUAAAU(A)DD substrate by the full CCR4–NOT complex. Reaction conditions are similar to (A) except full-length CSDE1 was used in these experiments. (C) Reduction of viral mRNA poly(A)-tail length in cells depleted of CSDE1. Shown is an RNase H northern blot measuring poly(A)-tail lengths of viral mRNAs in virions for each passage of a serial infection in control and CSDE1 KD NIH-3T3 cells. (D) CDF plot of the poly(A)-tail lengths of viral mRNAs in virions quantified from (C). Plotted are the average values of two independent nontargeting sgRNAs or two sgRNAs targeting *Csde1*.

We next returned to our original serial infection scheme (**Figure 1A**) and tested whether CSDE1 has a role in the poly(A)-tail length distribution of viral mRNAs in infected cells. To this end, we knocked down CSDE1 using CRISPRi with two independent guide RNAs (**Figure S4A**) and used an RNAse H northern blot to analyze the tail-length distribution of viral mRNAs at each passage of a 5-passage serial infection. Consistent with the in vitro results, infections in CSDE1 KD cells resulted in shorter poly(A)-tail lengths compared to infections in control cells (**Figures 4C and 4D**).

Interestingly, although the poly(A)-tail lengths of viral mRNAs were consistently shorter in CSDE1 KD cells, the extent of poly(A)-shortening was not exacerbated as infection passages increased, suggesting that a new steady-state is achieved during the first passage.

Taken together, our findings show that binding of CSDE1 to the tail-bound PABPC1 protects mRNAs from deadenylation, and loss of CSDE1 results in shortened poly(A) tails of viral mRNAs. Notably, because CSD2 can only interact with the RRM1 of a terminal PABPC1—its lateral face being inaccessible in non-terminal PABPC1 molecules^53^—this interaction occurs exclusively at the 3′ end of the poly(A) tail. We therefore refer to this assembly as the poly(A)-tail capping complex (PCC).

### PABPC1 activity is limiting during infection, which destabilizes short-tailed host mRNAs

Host mRNA degradation during coronaviral infection has been documented in multiple studies, with nsp1 thought to play a central role in this process^54–56^. However, for SARS-CoV-2, the correlation between mRNA half-life changes observed during infection and those observed under ectopic nsp1 expression is weak^54,57^ (*R*_s_ = 0.11, **Figure 5A**). For MHV, we first confirmed that overexpression of MHV nsp1, similar to overexpression of SARS-CoV-2 nsp1^58,59^, attenuates the amount of nascent peptides that can be labeled with puromycin—a proxy for global translation levels (**Figure 5B**)—and reduces cytoplasmic mRNA levels (**Figure 5C**). We then compared the changes in cytoplasmic mRNA abundance upon ectopic expression of nsp1 (6 and 12 h of expression) with those during MHV infection (9 and 11 hpi). As observed for SARS-CoV-2 nsp1, the effect of MHV nsp1 expression explained only a small portion of host mRNA degradation observed during infection (*R*_s_ = 0.14 and 0.25, **Figures 5D and 5E**, respectively). We also determined that RNase L activation was not contributing to mRNA degradation, as cells at the final timepoint of infection (i.e., 11 hpi) had intact rRNAs (**Figure S5A**). Taken together, these observations suggest the existence of one or more nsp1-independent mechanisms that contribute to host mRNA decay during coronaviral infection.

**Figure 5.**
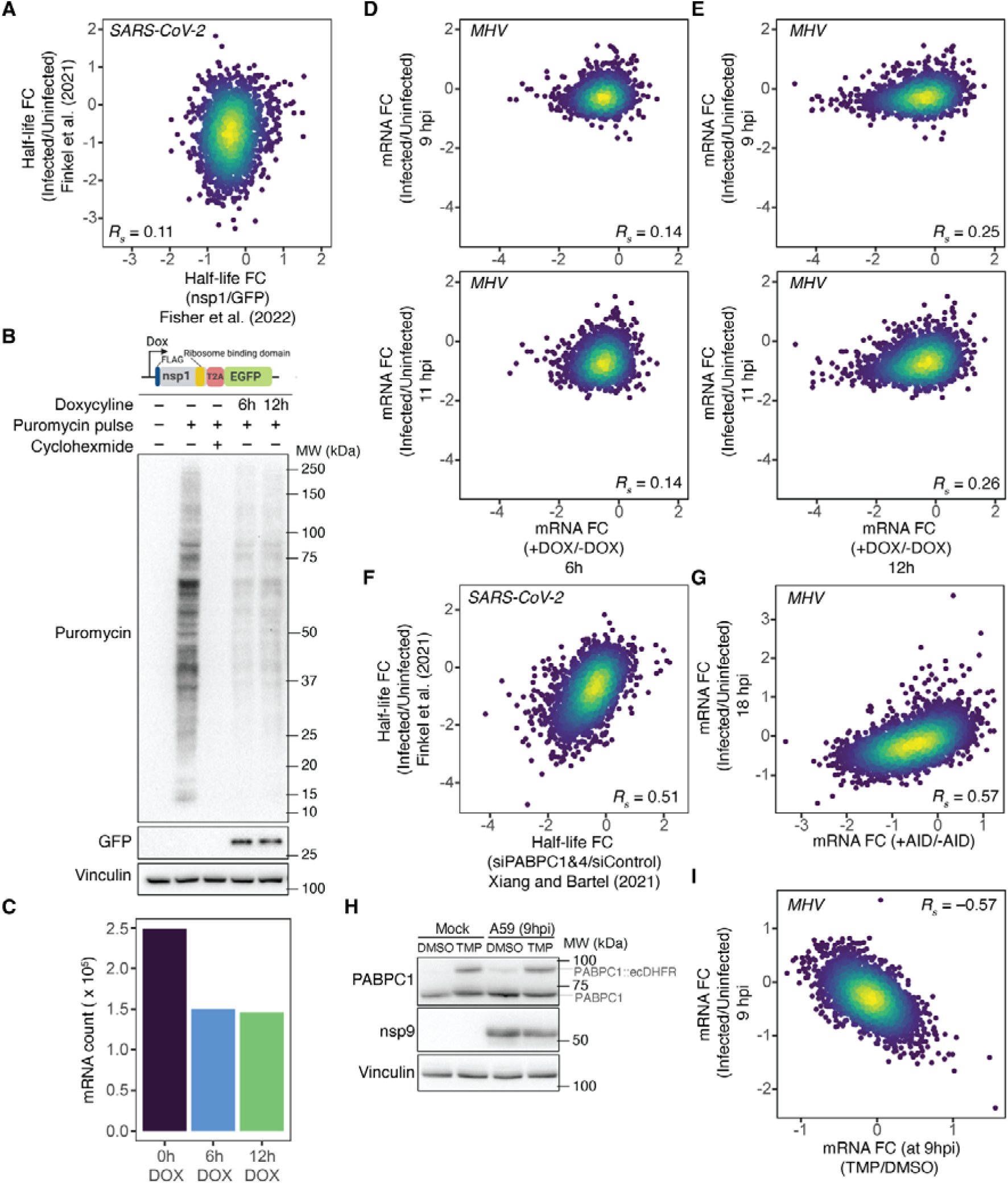
PABPC1 activity is limiting during infection, which destabilizes short-tailed host mRNAs. (A) A very weak relationship between changes in half-lives of host mRNAs observed upon infection with SARS-CoV-2 in Calu-3 cells^57^ and changes in half-lives of mRNAs observed upon expression of SARS-CoV-2 nsp1 in HEK-293T cells^54^. (B) Impact of MHV nsp1 on global translation levels. Shown are translating proteins labeled by puromycylation in L2 cells expressing a doxycycline-inducible version of MHV nsp1 for the indicated times. The lack of signal in the presence of cycloheximide is shown as a control. (C) Degradation of mRNAs by MHV nsp1. Plotted are abundances of cytoplasmic mRNAs in L2 cells expressing a doxycycline-inducible version of MHV nsp1 for the indicated time. (D) The correlation between changes in mRNA abundances in L2 cells infected with MHV for either 9 or 11 h (top and bottom, respectively) and changes in mRNA abundances of L2 cells expressing a dox-inducible version of nsp1 from MHV for 6 h. (E) The correlation between changes in mRNA abundances in L2 cells infected with MHV for either 9 or 11 h (top and bottom, respectively) and changes in mRNA abundances of L2 cells expressing a dox-inducible version of nsp1 from MHV for 12 h. (F) The correlation between changes in half-lives of host mRNAs upon infection with SARS-CoV-2 in Calu-3 cells^57^ and changes in half-lives of mRNAs upon knockdown of PABPC1 and PABPC4 in HeLa cells^11^. (G) The correlation between changes in mRNA abundances upon infection of HCT116 cells with MHV (18 hpi) and changes in mRNA abundances upon PABPC1 depletion in HCT116 cells. (H) Ectopic increase of PABPC1 levels during MHV-A59 infection. Shown are Immunoblots of L2 cells expressing a PABPC1 variant in which the C term of PABPC1 is fused to the destabilizing moiety ecDHFR in mock and infected cells treated with either DMSO or stabilizing ligand TMP. (I) Sensitivity of infection-induced cytoplasmic mRNA degradation to increased PABPC1 levels. Plotted is the correlation between changes in cytoplasmic host mRNA abundances of L2 cells upon infection with MHV and changes in cytoplasmic host mRNA abundances of MHV-infected L2 cells upon ectopic PABPC1 expression upon treatment with TMP.

Viral mRNAs maintain a poly(A)-tail length of 60–80 nt, with 60 nt being the footprint length for two closely spaced PABPCs^60,61^. This, combined with the cooperative nature of PABPC binding to the poly(A) tail^53,62^, prompted us to consider the possibility that maintaining this tail length allows the viral mRNAs to effectively compete with host mRNAs for PABPCs. One implication of this model, in which mRNAs compete for limiting PABPC, is that host mRNA degradation during infection might resemble that observed under limiting PABPCs levels.

To determine whether limiting PABPC1 contributes to host mRNA degradation during infection, we compared the changes in host mRNA half-lives during SARS-CoV-2 infection^57^ with changes observed upon knockdown of PABPC1 and PABPC4^11^. Because these measurements were performed in different cell lines—Calu-3 for SARS-CoV-2 infection and HeLa for PABPC knockdown—we first confirmed that mRNA half-lives were comparable between the two cell lines (**Figure S5B**). Despite some small differences between cell lines, we found a strong correlation between half-life changes observed during infection of Calu-3 cells and those observed upon PABPC1 and PABPC4 knockdown in HeLa cells (*R*_s_ = 0.51, **Figure 5F**). To avoid effects of long-term of PABPC knockdown^63^, and to investigate whether the correspondence we observed with SARS-CoV-2 infection also holds for cells infected with MHV, we utilized a cell line that uses an auxin-inducible degron (AID) system to rapidly deplete PABPC1^11^. Because these cells lack the receptor required for MHV infection, we expressed the MHV entry receptor, Cecam1^64,65^, in this cell line and compared changes in mRNA abundance between MHV-infected cells and cells depleted of PABPC1 using the AID system. Consistent with the SARS-CoV-2 findings, we found that changes in mRNA abundance observed upon MHV infection strongly correlated with those observed upon PABPC1 depletion (*R*_s_ = 0.57, **Figure 5G**).

To further validate that PABPC limitation during infection contributes to host mRNA degradation, we temporally increased PABPC1 levels during infection by stabilizing a PABPC1 construct containing the destabilizing degron ecDHFR^66^. Under DMSO-treated conditions, this construct is rapidly degraded; however, upon treatment with the ecDHFR ligand Trimethoprim (TMP), the PABPC1-ecDHFR is stabilized, thereby increasing the total level of PABPC1. Using this setup, we infected cells with MHV, treated them with TMP at 5 hpi, harvested the cells at 9 hpi, and analyzed changes in cytoplasmic mRNA abundance via RNA-seq (**Figure 5H**). Changes in mRNA abundance observed during infection strongly corresponded to those observed upon TMP treatment of infected cells (*R*_s_ = –0.57, **Figure 5I**), which indicated that the temporal overexpression of PABPC1 can reverse the reduction of cytoplasmic host mRNAs during infection, as expected if limiting PABPC activity causes degradation of host mRNA during infection.

### Host mRNAs with short poly(A) tails are preferentially degraded during infection

When PABPC activity becomes limiting in post-embryonic cells, mRNAs with short tails are preferentially destabilized^11^ (**Figure S6C and S6D**). Therefore, to examine further the hypothesis that limiting PABPC was affecting host mRNA stability, we looked for evidence of loss of short-tailed mRNAs in infected cells. Interestingly, this analysis revealed a preferential loss of host mRNAs with shorter tail lengths as infection progressed (**Figures 6A and 6B**), which was not observed in our spiked-in zebrafish control mRNAs (**Figure S6A**). Notably, the strongest depletion of short-tailed mRNAs occurred at 9 and 11 hpi, coinciding with the times at which viral mRNA levels matched and exceeded the abundance of cytoplasmic host mRNAs (**Figure S6B**). A loss of short-tailed mRNAs was also observed when analyzing the average behavior of the hundreds of different mRNAs that passed our expression cutoffs, which showed that this signature feature of limiting PABPC activity was occurring generally, not just for a small number of highly expressed mRNAs (**Figure 6C**).

**Figure 6.**
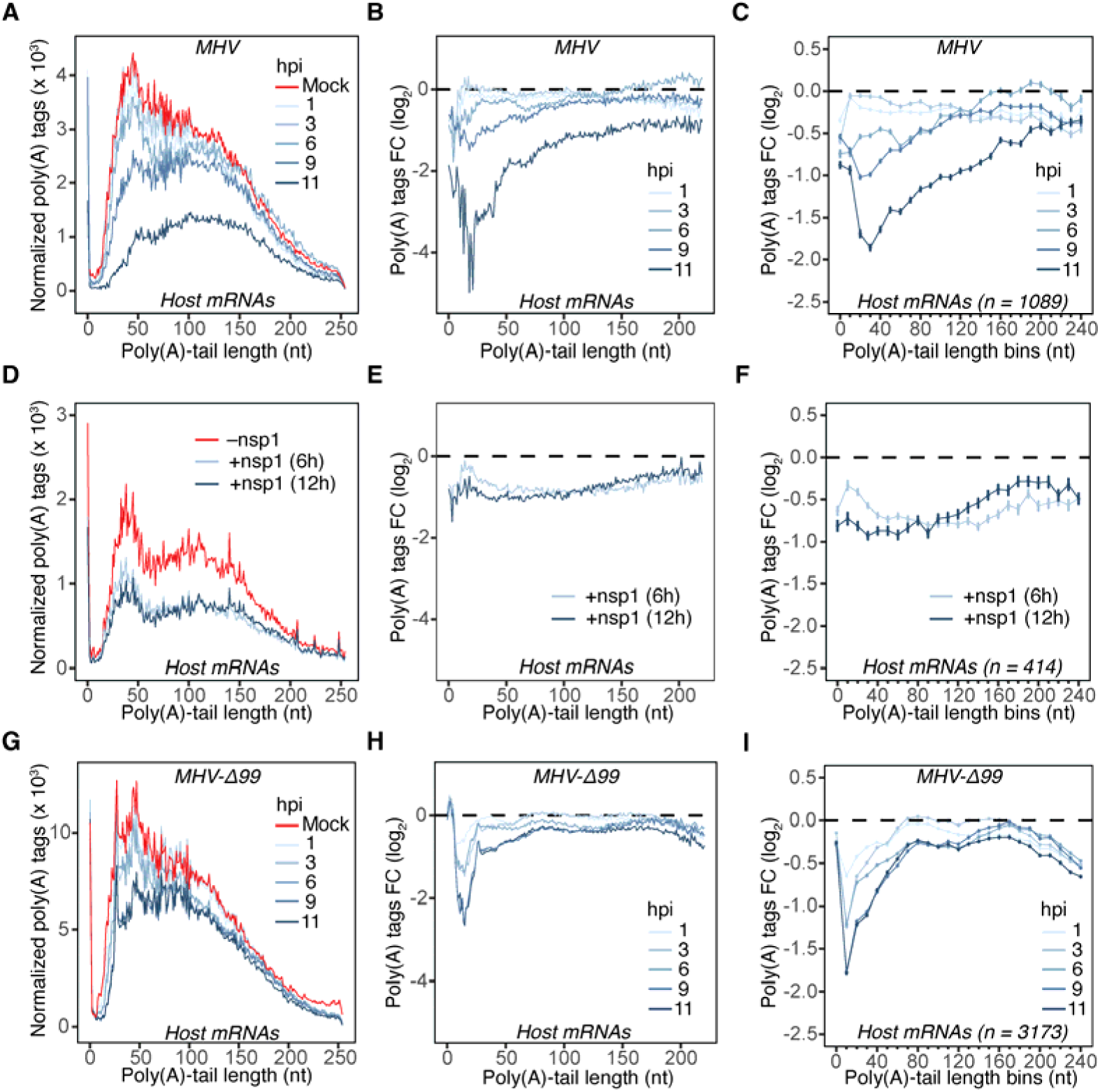
Host mRNAs with short poly(A) tails are preferentially degraded during infection. (A) Preferential degradation of cytoplasmic short-tailed host mRNAs during infection with MHV-A59. Plotted are normalized tail-length abundances of host mRNAs in mock and MHV-infected L2 cells. (B) Preferential degradation of cytoplasmic short-tailed host mRNAs during infection with MHV-A59. Plotted are changes in abundance of host mRNAs of the indicated tail lengths observed upon infection. (C) Preferential degradation of cytoplasmic short-tailed host mRNAs during infection with MHV-A59, analyzed after grouping mRNAs by gene. Plotted for each tail-length bin is the average change observed per gene. (D) Tail-length-agnostic degradation of cytoplasmic mRNAs upon expression of MHV nsp1. Plotted are normalized tail-length abundances of mRNAs in L2 cells expressing dox-inducible version of MHV nsp1 as well as noninduced cells. (E) Tail-length-agnostic degradation of cytoplasmic mRNAs upon expression of MHV nsp1. Plotted are changes in abundance of host mRNAs of the indicated tail lengths observed upon expression of MHV nsp1. (F) Tail-length-agnostic degradation of cytoplasmic mRNAs upon expression of MHV nsp1, analyzed after grouping mRNAs by gene. Plotted for each tail-length bin is the average change observed per gene. (G–I) Preferential degradation of cytoplasmic short-tailed host mRNAs during infection with MHV-Δ99; otherwise as in (A), (B), and (C), respectively.

We next asked to what extent expression of nsp1 can explain the bias in degradation of short-tailed mRNAs. Ectopic expression of MHV nsp1 (6 and 12 hours) did not lead to preferential degradation of short-tailed mRNAs (**Figures 6D–F and Figure S6E**). When considering that the nsp1-mediated degradation of mRNAs occurs through the nsp1 binding of the host ribosome^55^, our observation is consistent with the absence of coupling between poly(A)-tail length and translation efficiency in post-embryonic cells^10^. To explore further the potential contribution of nsp1, we utilized an nsp1-mutant MHV strain, MHV-Δ99^67^, in which a substantial portion of the C-terminal helical bundle responsible for 40S ribosomal subunit binding is deleted. We first confirmed that the nsp1 from MHV-Δ99 is indeed defective in attenuating global translation levels compared to wildtype nsp1, using puromycylation of nascent peptides as a readout (**Figures S6F and S6G**). We then repeated our infection time course using the mutant virus (**Figure S2A**) and examined the poly(A)-tail length distributions. Remarkably, although the extent of host mRNA degradation was attenuated in MHV-Δ99, as might be expected for a hypomorphic viral mutant, we still observed preferential loss of short-tailed host mRNAs (**Figures 6G-6I** and **Figure S6H**). These observations strongly support an nsp1-independent mode of host mRNA degradation during infection.

Taken together, our findings support a model in which during the course of infection, PABPC activity becomes limiting, such that viral mRNAs compete with host mRNAs for PABPC binding. In this regime in which short-tailed mRNAs are destabilized, maintaining the poly(A)-tail length of viral mRNAs at a competitive length for binding to PABPC1 provides a benefit that helps the virus dominate the cytoplasm of the infected cells.

## Discussion

RNA viruses exhibit striking diversity in how they configure the 3′ ends of their genomes and transcripts, reflecting distinct solutions to the universal challenges of RNA stability, translation, and replication. Flaviviruses, for instance, encode structured RNA elements such as stem–loops at their 3′ ends, whereas picornaviruses and coronaviruses use poly(A) tails that mimic those of host mRNAs^68^. In polyadenylated RNA viruses that do not replicate through a DNA intermediate, the poly(A) tail is templated from a poly(U) segment in the negative-strand intermediate. This tail protects genomic RNA from 3′ trimming and delays entry into the mRNA-decay pathways triggered once tails shorten below ∼25 nucleotides^4^. Because these viruses lack access to nuclear polyadenylation machinery, mechanisms that restore or preserve tail length in the cytoplasm must be employed.

Our findings reveal two complementary mechanisms coronaviruses use to protect their poly(A) tails (**Figure 7**). First, cytoplasmic polyadenylation of newly transcribed viral mRNAs occurs within double-membrane vesicles (DMVs) derived from the ER. Thus, DMVs not only shield viral dsRNA intermediates from host sensors, but also provide a protected environment for the synthesis of long poly(A) tails, insulating them from cytoplasmic deadenylation machinery. More broadly, our results add to emerging evidence—including cytoplasmic polyadenylation that can add mixed tails (which include a few non-A nucleotides) in other settings^69–73^—that deadenylation is not necessarily irreversible. Such post-embryonic cytoplasmic polyadenylation has been reported primarily for viral RNAs and, to date, has only rarely been observed for nuclear-encoded mRNAs^74–76^.

**Figure 7.**
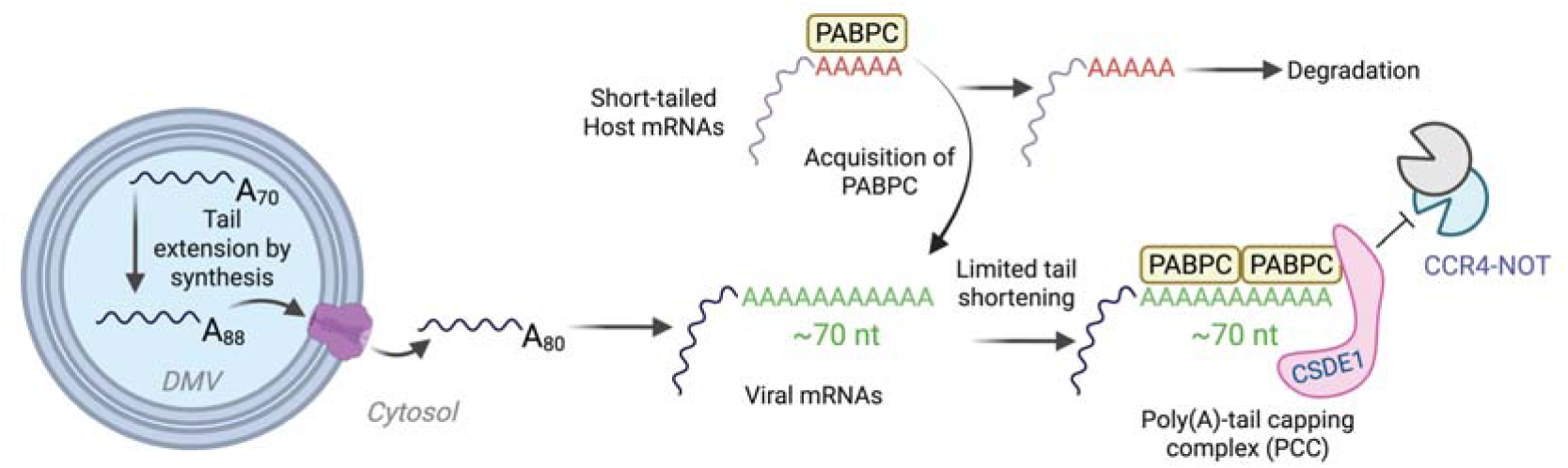
Model for viral strategies to preserve poly(A)-tail length and promote degradation of short-tailed host mRNAs during infection. Coronaviruses employ two complementary mechanisms to safeguard the poly(A) tails of their mRNAs: (1) nontemplated tail extension during synthesis within the protected environment of double-membrane vesicles (DMVs), and (2) repression of deadenylation by the poly(A)-tail capping complex (PCC). By preserving tails long enough to bind at least two PABPC molecules, viral mRNAs secure preferential access to limiting PABPC, thereby depriving short-tailed host mRNAs of PABPC binding and promoting their degradation.

Second, we identify the PCC, consisting of the poly(A) tail, PABPC1, and the pro-viral protein CSDE1. Although CSDE1–PABPC1 interaction has been reported before^51,52^, our study advances the understanding of this interaction in three ways: (1) we map the interaction to the CSD2 domain of CSDE1 and the RRM1 domain of PABPC1, (2) we show that CSDE1, like eIF4G^77^, preferentially binds poly(A)-bound PABPC1, and (3) we demonstrate that the poly(A) tail is more resistant to deadenylation by CCR4–NOT when CSDE1 is bound to PABPC1–poly(A) complex compared to PABPC1–poly(A) alone. A key open question is whether viral mRNAs contain sequence elements that that help recruit CSDE1. Such elements would enable viral mRNAs to preferentially benefit from CSDE1-mediated protection. Seven of nine CSDs in CSDE1 are annotated as RNA-binding domains; these provide a potential starting point for investigating whether interactions with viral RNA might impart specificity to CSDE1 association.

Viral mRNAs appear to dominate the host cytoplasm (**Figure S6B**) with poly(A) tails long enough to accommodate at least two PABPC molecules (> 50 nt). Previous studies have described the dual roles of PABPC in regulating translation efficiency in oocytes and early embryos, and stabilizing mRNA in post-embryonic cells^11,78^. In the latter case, when cytoplasmic levels of PABPC are reduced to become limiting relative to the abundance of PABPC binding sites on poly(A) tails, short-tailed mRNAs compete poorly for the remaining PABPC and are degraded, presumably through decapping of mRNAs lacking bound PABPC^11,79^. Viral infection creates an interesting scenario in which, instead of changes in PABPC levels, the abundance of PABPC binding sites increases due to cytoplasmic transcription of polyadenylated viral mRNAs. In this study, we show that coronavirus infection creates an environment in which short-tailed mRNAs are preferentially degraded. Indeed, depletion of short-tailed mRNAs from the cytoplasm explains why we and others have observed an increase in the median tail length of host mRNAs increases during coronaviral infection^80^. Importantly, we also demonstrate that increasing PABPC1 levels during infection rescues the degradation of host mRNAs, consistent with additional PABPC1 becoming available to bind short poly(A)-tailed transcripts. Moreover, CSDE1 might provide viral mRNAs with an additional advantage in acquiring PABPC, beyond the inherent effect of tail length.

We show that depletion of short-tailed host mRNAs during infection operates in parallel with nsp1 to induce degradation of host mRNAs. An intriguing possibility is that limiting PABPC enables the virus to use preferential PABPC binding to its own mRNAs to promote degradation of short-tailed isoforms of pre-existing host mRNAs, while nsp1-mediated degradation neutralizes newly transcribed host mRNAs, which have longer poly(A)-tail lengths and include those transcribed in response to infection. This is consistent with proposed role of nsp1 in counteracting the interferon response, which is activated in response to infection^54^.

Strategies to preserve cytoplasmic poly(A) tails might also exist beyond coronaviruses. Previous studies have described CSDE1 as a pro-viral factor for other viruses such rhinovirus and poliovirus, which similarly to coronaviruses are cytoplasmic RNA viruses with polyadenylated genomes. For these viruses, CSDE1 has been shown to be an internal ribosome entry site (IRES) trans-activating factor (ITAF) by binding the type I IRES in 5′ UTR of the viral genomic mRNA and promoting translation initiation^81–84^. Coupling this interaction with the IRES with its interaction with PABPC1, CSDE1 might act to circularize the genome^85^ by simultaneously bridging the 5′ and 3′ ends of viral RNAs, while protecting the poly(A) tails against deadenylation. Moreover, the LINE-1 retrotransposon encodes a PABPC interacting and essential element (PIE) that is predicted by Alphafold to target the same lateral face of PABPC1 RRM1^86^ as CSDE1, and the LINE-1 poly(A) tail and its length are critical determinants of retrotransposition efficiency^87–89^. Expanding the search for PABPC1 interactors, followed by functional validation, might reveal additional RRM1-binding proteins that influence deadenylation, uncovering yet more viral and cellular strategies to stabilize poly(A) tails.

## Acknowledgements

We thank current and former members of the Bartel lab—especially Thy Pham, Charlie Shi, Lianne Blodgett, Kehui Xiang, Sean McGeary, Daniel Lin, and Jordan Ray—as well as Jimmy Ly and Iain Cheeseman for their helpful discussions and insights. The MHV-Δ2-GFP3 viral strain was a generous gift by Mark Denison and MHV-Δ99 was kindly provided by Volker Thiel. The L2 cell line was a generous gift from Susan Weiss. We also thank Hans-Heinrich Hoffmann and Charlie Rice for providing us with total RNA collected from SARS-CoV-2 infected Huh-7 cells. This research was supported by the Intramural Research Program of the National Institutes of Health (NIH). The contributions of the NIH authors are considered Works of the United States Government. The findings and conclusions presented in this paper are those of the authors and do not necessarily reflect the views of the NIH or the U.S. Department of Health and Human Services. Funding for this study was provided by the National Institutes of Health, National Cancer Institute (K00CA234921 to A.L.), and Intramural Research Program of the National Institutes of Health, National Cancer Institute (ZIABC011977 to E.V.). D.P.B. is an investigator of the Howard Hughes Medical Institute. We thank Sumeet Gupta, Amanda Chikala, and Stephen Mraz from the Whitehead Institute Genome Technology Core for assistance with sequencing.

## Author Contribution

A.L and D.P.B. conceived the project, design the study, and wrote the manuscript. A.L. conducted majority of experiments with help from S.S.N. for data analysis. In vitro deadenylation experiments as well as related protein purifications were performed by Y.L. and supervised by E.V.. A.B. conducted the proteome-wide ColabFold screen with RRM domains and CSD2.

## Material and Methods

### Cell culture

The L2 cell line (from Susan Weiss) was maintained at 37°C with 5% CO_2_ in DME10 media which consists of Dulbecco Modified Eagle Medium (DMEM, Corning) supplemented with 10% Fetal Bovine Serum (FBS, Takara), 0.37% sodium bicarbonate, 2 mM L-glutamine and 2.5 µg/mL of Amphotericin B (Thermo Fisher) at 37°C incubator with 5% CO_2_. HCT116 PABPC1-AID were obtained from Kehui Xiang and cultured in McCoy’s 5A media (Thermo Fisher) supplemented with 10% FBS and 2 mM L-glutamine. NIH-3T3 cells expressing dCas9 were obtained from the Weissman lab and cultured in DMEM supplemented with 10% BCS (Sigma). HEK-293T cells were obtained from ATCC and cultured in DMEM supplemented with 10% FBS.

### Plasmids

All DNA plasmids were assembled using HiFi DNA Assembly (NEB). Plasmids and their sequences information will be available at Addgene.

### Viral Strains and propagation

The MHV-A59 strain was obtained from ATCC, the MHV-Δ2-GFP3 strain was a generous gift by Mark Denison, and MHV-Δ99 was kindly provided by Volker Thiel. Upon receipt, a plaque assay was conducted on each virus, and viruses were plaque purified for downstream experiments.

For propagation, L2 cell monolayers were infected at a low multiplicity of infection (0.1 MOI) and cultured until extensive cytopathic effect was observed (∼ 2-3 days). Infected cells were harvested by a single freeze–thaw cycle, followed by sonication on ice to release intracellular virus (three 20-second bursts at 100 W with cooling intervals), and clarification by centrifugation at 1,500 x g for 10 min at 4°C. Supernatants were aliquoted and stored at −80 °C as working stocks and viral titer was determined prior to downstream experiments.

### Plaque assay and purification

Infectious virus titers were determined by plaque assay using L2 cells as described in Leibowitz et al.^90^. Briefly, confluent L2 monolayers in 6-well plates were washed with serum-free medium and inoculated with 0.5 mL of 10-fold serial dilutions of virus in duplicate. After 1 h of adsorption at room temperature with gentle rocking, cells were overlaid with a 1:1 mixture of 1.6% agarose and 2× DMEM containing 2% serum and incubated at 37 °C and 5% CO. Plaques were visible after 2-3 days and were visualized by crystal violet staining following removal of the agarose overlay. For plaque purification, prior to staining, visible well-isolated plaques were picked with a sterile Pasteur pipette, resuspended in medium, and used to infect L2 cells for preparation of low-passage seed stocks.

### Infection

All infections were conducted at an MOI of 5 unless noted otherwise. Infection with MHV strains were performed according to Leibowitz et al.^90^. Briefly, cells were removed from complete media, washed once with serum free media, and then incubated the inoculum in DME2 (otherwise same as DME10 but with 2% FBS) with gentle mixing at room temperature for one hour. The inoculum was then removed and the cells were washed once with serum free media and then incubated with DME2 at 37°C incubator and harvested later at indicated times. The percentage of infected cells was determined by development of the cytopathic effect or by immunofluorescence staining of viral protein nsp9 in parallel.

### Serial Infection

In serial infection experiments, successive infections were carried out in 15 cm dishes, where for each passage, the conditioned media was collected at 12 hours post infection and virions were pelleted as described in Leibowitz et al.^90^ and split in half for RNA collection or subsequent infection. The viral titer for each round was determined using plaque assay and MOI was kept the same throughout infection rounds.

### Subcellular Fractionation

Subcellular fractionation was carried out as described previously with minor modifications^57^. Mock- or MHV-infected cells grown in 15-cm dishes were washed twice with ice-cold PBS and collected by scraping. Ten percent of the cells were aliquoted as input, and the remaining cells were pelleted at 300 × g for 5 min. The pellet was resuspended in 150 µL of buffer A (15 mM Tris-Cl pH 8, 15 mM NaCl, 60 mM KCl, 1 mM EDTA pH 8, 0.5 mM EGTA pH 8, 0.5 mM spermidine, 10 U ml ¹ RNase inhibitor) and then gently mixed with 150 µL of 2× cytoplasmic lysis buffer (15 mM Tris-Cl pH 8, 15 mM NaCl, 60 mM KCl, 1 mM EDTA pH 8, 0.5 mM EGTA pH 8, 0.5 mM spermidine, 10 U ml ¹ RNase inhibitor, 0.5% NP-40, and 2× Halt protease inhibitor) and incubated on ice for 10 min. After centrifugation at 400 × g for 5 min, the supernatant (cytoplasmic fraction) was collected and clarified by an additional spin at 500 × g for 1 min, then divided for RNA extraction (TRIzol) and protein analysis. The nuclear pellet was resuspended in 1 mL RLN buffer (50 mM Tris-Cl pH 8, 140 mM NaCl, 1.5 mM MgCl, 0.5% NP-40, 10 mM EDTA, 10 U ml ¹ RNase inhibitor), incubated on ice for 5 min, and centrifuged at 500 × g for 5 min. The final nuclear pellet was processed for RNA extraction with TRIzol or protein analysis using RIPA buffer (20 mM Tris-HCl pH 8.0, 150 mM NaCl, 60 mM KCl, 1 mM EDTA, 1 mM EGTA, 0.5 mM spermidine, 1% NP-40, 1% sodium deoxycholate, 1× Halt protease inhibitor).

### DMV immunoprecipitation

DMV immunoprecipitations were adapted from prior studies with modifications^27,91^. Infections were initiated in two 500-cm² plates, and at 11hpi cells were washed twice with ice-cold PBS, scraped, and pelleted at 300 × g for 5 min. Pellets were resuspended in KPBS buffer and gently dounce homogenized with 20 strokes. The homogenate was clarified at 1000 × g for 5 min, and the supernatant was subjected to GFP immunoprecipitation using GFP-Trap beads (Proteintech) for 30 min at 4 °C. Beads were washed three times with KPBS, with transfers to new tubes during the first and final washes. For RNA analysis, RNA was extracted directly from beads using TRIzol, whereas for protein analysis, beads were boiled in 1× LDS sample buffer (Invitrogen).

### Nascent RNA labeling

To label nascent transcripts, infected cells were incubated with 200 µM 5-ethynyl uridine (5-EU; Jena Biosciences) for 10 min. Following labeling, cells were washed once with ice-cold PBS, scraped, and pelleted by centrifugation at 4 °C. The cytoplasmic fraction was then isolated, and RNA was extracted from this fraction using Trizol. The click-chemistry and streptavidin capture of 5-EU labeled RNAs were carried out as described previously^4^, with minor modifications.

#### Click-chemistry biotinylation

Reactions were assembled in 100 µL containing 250 µg or 100 µg of total RNA in 50 mM HEPES (pH 7.5), 4 mM disulfide-biotin azide (Vector Labs), 2.5 mM CuSO (MilliporeSigma), 2.5 mM THPTA (Vector Labs), 10 mM sodium ascorbate (MilliporeSigma), and 5 ng of standard GFP mRNA labeled with 5-EU (internal control). Reactions were incubated at room temperature for 1 h protected from light, then quenched with 5 mM EDTA. RNA was extracted with phenol:chloroform (pH 8.0), precipitated with linear acrylamide and ethanol, and resuspended in 100 µL of 1× HSWB buffer (10 mM Tris pH 7.4, 1 mM EDTA, 100 mM NaCl, 0.01% Tween-20).

#### Streptavidin pulldown

Biotinylated RNAs were captured using MyOne C1 streptavidin beads (Thermo Fisher; 100 µL beads per 25 µg of labeled RNA) pre-washed twice each with 1 mL of 1× B&W buffer (10 mM Tris pH 7.4, 1 mM EDTA, 100 mM NaCl), 1 mL of Solution A (0.1 M NaOH, 50 mM NaCl), 1 mL of Solution B (0.1 M NaCl), and 1 mL of water. Beads were blocked with 0.5 µg/µL yeast total RNA in 1 mL of 1× HSWB buffer for 30 min at 23 °C with intermittent mixing.

Blocked beads were incubated with the 100 µL of biotinylated RNA from the previous step for 30 min at 23 °C, washed three times with 1× HSWB, then washed twice with pre-warmed (50 °C) water and twice with pre-warmed (50 °C) 10× HSWB. Biotinylated RNAs were eluted by reducing the disulfide linkage with 180 µL of 0.5 M TCEP for 20 min at 50 °C with mixing. The eluate was collected, combined with a rinsing the beads with 150 µL of water, and ethanol-precipitated in the presence of 0.3 M NaCl and linear acrylamide. The recovered RNA was washed with 70% ethanol and resuspended in nuclease-free water for downstream library preparation.

#### In vitro transcription of standard 5-EU labeled GFP mRNA

5-EU labeled GFP mRNAs were generated by in vitro transcription of PCR-amplified templates encoding GFP with a T7 promoter using the MEGAscript T7 Transcription Kit (Thermo Fisher). 20 µL reactions containing 0.1 µM of template, 2 µL each of 100 mM ATP, GTP, and CTP, 2 µL of 100 mM UTP, 1 µL of 10 mM 5-ethynyl-UTP (Jena Biosciences; corresponding to a 20:1 ratio of UTP:5-EUTP), 2 µL of 10× T7 reaction buffer, 2 µL of T7 enzyme mix, and 1 µL of 0.1 M DTT. Reactions were incubated at 37 °C for 2 h, followed by DNase treatment according to the manufacturer’s instructions. The transcribed RNAs were gel purified and eluted overnight at 22 °C in elution buffer (10 mM HEPES pH 7.5, 0.3 M NaCl). Eluted RNAs were ethanol-precipitated, resuspended in nuclease-free water, and stored at –80 °C.

### Puromycylation

For global analysis of protein translation, nascent chains were label with puromycin, by treating the cells with 1 µg/ml of puromycin (Thermo Fisher), and cells were maintained at 37 °C for 10 minutes before collection. In parallel control samples, translation was blocked by adding cycloheximide (100 µg/ml) 5 minutes prior to puromycin treatment.

### RNase H Northern Blot

RNA from virions collected from conditioned media from each infection round during serial infection was used for RNase H Northern blot as described previously^11^. The following are sequences of the oligo used for generating 3_′_-end fragments as well as the probe used for northern blotting

3′-end RNase H oligo: AGGAATAGTACCCTGATGTGAGCTCTTCCC

Northern blot oligo: GATTCTTCCAATTGGCCATGATCAACTTCATTCATTTACTAGGGCAT

### Immunoblot

Protein samples for western blotting were prepared by boiling the lysates in NuPAGE LDS Sample Buffer (Thermo Fisher, #NP0007) diluted to 1× and supplemented with 1 mM DTT. Proteins were separated on NuPAGE 4–12% Bis-Tris 1.0 mm gels (Thermo Fisher, #NP0321BOX) and transferred to 0.45 µm PVDF membranes (Thermo Fisher, #88518) using a mini gel tank (Thermo Fisher, #A25977). Electrophoresis was performed at 170 V for 56 min, and transfers were carried out at 30 V for 60 min. Following transfer, membranes were blocked in TBS containing 0.05% Tween-20 (TBST) and 10% BSA for 1 h at room temperature, then incubated overnight at 4 °C with primary antibodies diluted in the same buffer. The primary antibodies were used at the following dilutions: PABPC1 (Cell Signaling Technology, #4992S, 1:1000), Histone H3 (Cell Signaling Technology, #9715S, 1:5000), nsp9 (MHV; Thermo Fisher, #200-301-A56, 1:1000), LC3 (Cell Signaling Technology, #2775S, 1:1000), CALR (Cell Signaling Technology, #12238S, 1:1000), CTSB (Cell Signaling Technology, #31718S, 1:1000), Golgin-97 (Cell Signaling Technology, #13192S, 1:1000), CSDE1 (Abcam, #ab201688, 1:1000), Vinculin (Cell Signaling Technology, #13901S, 1:5000), GFP (Cell Signaling Technology, #2956S, 1:2000), and Puromycin (MilliporeSigma, #MABE343, 1:1000). Membranes were washed three times for 10 min each with TBST, incubated with secondary antibody for 1 h at room temperature, and washed again three times for 10 min each in the same buffer. The membranes were then imaged using ChemiDoc™ MP Imaging System (Biorad).

### CRISRPi

NIH-3T3 cells expressing dcas9-GFP were used for CRISPRi experiments. Specifically, plasmids containing sgRNAs were cloned in pLG1 plasmid (Addgene) by restriction cloning. Lentivirus was produced in HEK-293T cells cultured in 6-well plates. For each well, cells were transfected with 1.4 µg of sgRNA plasmid, 0.94 µg of dR8.91 packaging plasmid (Addgene), and 0.47 µg of pMD2.G envelope plasmid (Addgene) using FuGENE 6 and Opti-MEM. After 16 h, the medium was replaced, and 48 h later, the viral supernatant was collected and centrifuged at 500 × g for 10 min to remove debris. To infect NIH-3T3 cells cultured in 6-well plates, 500 µL of virus-containing supernatant (approximately 40% of the total) was mixed with 1 mL of fresh medium and polybrene to a final concentration of 8 µg/mL. Cells were spinfected at 1,200 × g for 1.5 h at 22 °C and then incubated at 37 °C. The following day, cells were transferred to 10-cm dishes and cultured in medium containing 1 µg/mL puromycin. Selection continued for 3 days before cells were expanded into 15-cm dishes for serial infection experiments. The following are the sequences of sgRNAs used for knockdown experiments:

*Csde1* sgRNA-1: GTATGGCGGCGCTGGAGAGG

*Csde1* sgRNA-2: GCGGGCCGTGCTGCTTATGG

Non-targeting sgRNA-1: GGGAACCACATGGAATTCGA

Non-targeting sgRNA-2: GAGGTTACCCACCCAGCGGT

### Immunofluorescence

Cells grown on glass coverslips were infected with MHV-Δ2-GFP3 for 9 hours, then fixed and permeabilized with methanol. The slides were then blocked with 10% bovine serum albumin diluted in PBS. For staining of nsp9, the cells were incubated for 90 min with 1:500 dilution of nsp9 antibody (Thermo Fisher). The secondary staining was conducted for 1 hour with anti-mouse IgG-Alexa 568 Conjugate antibody (1:400, Thermo Fisher) and NucBlue (ThermoFisher) to label the nuclei. The cells were visualized with Super Resolution Structured Illumination Microscopy using a Zeiss LSM 980 with Airyscan 2 Laser Scanning Confocal with a 63x oil objective lens (W.M. Keck Biological Imaging Facility, Whitehead Institute). Image processing was performed with ImageJ software.

### Alphafold

#### Viral mRNA binding proteins and PABPC1

AlphaFold3 was used to predict multimeric complexes between full-length virus-associated RBPs and either full-length PABPC1 or its individual domains. The virus-associated RBPs were selected based on prior studies that identified proteins directly binding both genomic and subgenomic SARS-CoV-2 mRNAs^35,36^. A shared set of 96 candidate RBPs (‘baits’) was compiled from factors reported as significant in both studies. The PEAK score^40^ was then adapted to rank the predictions.

#### RRM domains and CSD2 domain of CSDE1

RRM domains from the human RBPome were extracted from RBPWorld^92^, a comprehensive database of RNA-binding proteins and domains derived from published experimental datasets and bioinformatic predictions. RNA-binding domains (RBDs) in the dataset were mapped to their corresponding Pfam IDs to query the Ensembl. For each gene, the canonical protein ID (ENSP) was retrieved along with the amino acid sequence, protein length, and Pfam domain coordinates. The curated list was then filtered for RNA recognition motifs, and the corresponding RRM domain sequences were extracted with 50 amino acids flanking each side or extended to the start/stop codons when the domain was near a boundary. The AlphaFold analysis for predicting multimers between fetched RRM domains (n= 350) and the CSD2 domain of CSDE1 was then performed using ColabFold^50^ followed by PEAK score^40^ to rank the predictions.

### Protein Purification

Reconstitution of the human CCR4–NOT complex^93,94^ and purification of PABPC1^95^ were described previously.

#### Full-length CSDE1

Full-length human CSDE1 carrying N-terminal His_6_-SUMO-tag was produced in BL21(DE3) Star *E.coli* cells using 2 L of autoinduction media at 20 °C overnight. The cells were harvested and resuspended in a lysis buffer containing 50 mM potassium phosphate, pH 7.5, 1 M NaCl, 25 mM imidazole, and then lysed by sonication. The lysate was clarified by centrifugation at 40,000 g for 45 min and loaded on a 5 mL Ni-charged HisTrap Excel column (Cytiva). The bound protein was washed with sixteen column volumes of lysis buffer and then with four column volumes of the same buffer containing 300 mM NaCl. The last round of wash was done with 2 column volumes of buffer containing 50 mM potassium phosphate, pH 7.5, 300 mM NaCl, 35 mM imidazole. His_6_-SUMO-CSDE1 was eluted from the column in 1 mL fractions with the second wash buffer supplemented with 250 mM imidazole. Peak fractions from Ni affinity chromatography were combined and incubated with HRV-3Cpro at 4 °C overnight to cleave off the His_6_-SUMO-tag. The protein mixture was then loaded and eluted on Superdex 200 26/600 (Cytiva) equilibrated in a buffer containing 10 mM HEPES/NaOH, pH 7.5, 200 mM NaCl, 5% (v/v) glycerol, 2 mM DTT. The peak fractions were then pooled together, concentrated to ∼5 mg/ml, flash-frozen in liquid nitrogen, and stored at -80 °C. The CSDE1^V136R, Y138R^ mutant was produced and purified identically to the wild-type version.

#### CSDE1 (2-186)

The production of CSDE1 (2-186) carrying an N-terminal His_6_-SUMO-tag was identical to that of the full-length version of CSDE1. Ni-affinity chromatography and cleavage by HRV-3Cpro were also performed identically to the full-length protein. To separate CSDE1 (2-186) from the cleaved His_6_-SUMO-tag and HRV-3Cpro, the protein mixture was diluted 1:1 with 50 mM HEPES/NaOH, pH 7.5 and applied to a 5 mL HiTrapQ column (Cytiva) equilibrated in 50 mM HEPES/NaOH, pH 7.5, 100 mM NaCl, 2 mM DTT. Purified CSDE1 (2-186) was captured in the flow-through fraction. Subsequently, it was loaded and eluted on Superdex 75 26/600 (Cytiva), equilibrated in a buffer containing 10 mM HEPES/NaOH, pH 7.5, 200 mM NaCl, 5% (v/v) Glycerol, 2 mM DTT. The peak fractions were then pooled together, concentrated to ∼5 mg/ml, flash-frozen in liquid nitrogen, and stored at -80 °C. CSDE1 (2-186)^V136R, Y138R^ mutant was produced and purified identically to the wild-type version.

### Size-exclusion chromatography

Formation of RNA–protein complexes was examined by size-exclusion chromatography (SEC) in 20 mM Tris (pH 8.0), 100 mM NaCl, 5% glycerol, and 2 mM DTT. Oligo(A)_10_ and oligo(A)_30_ (IDT) were gel purified prior to use. Samples were mixed and incubated on ice for 30 min, clarified by centrifugation at 17,000 × g for 5 min, and the supernatant was loaded onto the column. For minimal domains of CSDE1 and PABPC1, samples were run on a Superdex 200 Increase 10/300 GL column (Cytiva), and absorbance was monitored at 260 nm and 280 nm. For full-length proteins, samples were run on a Superdex 200 Increase 3.2/300 column, and absorbance was monitored at 220 nm. For Figures 3G and S3G, mixtures (total volume 500 µL) were prepared using 6 µM CSDE1 (2-186), 16 µM oligo(A)_10_, and 12 µM PABPC1 (2-191). For Figure S3C, mixtures (total 50 µL) were prepared using 375 nM full-length CSDE1, 750 nM full-length PABPC1, and 1.5 µM of oligo(A)_30._ For Figure S3D, mixtures (total 50 µL) were prepared using 375 nM full-length CSDE1, 750 nM full-length PABPC1, and 750 nM of oligo(A)_30._

### Mass photometry

Mass photometry measurements were carried out on Refeyn TwoMP instrument at Biophysical Instrumentation Facility at MIT. Prior to each experiment, the instrument stage and objective were cleaned, and a fresh coverslip was sequentially rinsed in isopropanol and ethanol and then dried with compressed air. The gasket was washed thoroughly with soap and water, rinsed, and dried prior to use. A drop of immersion oil was applied to the objective, and the coverslip and gasket were placed to form a sealed chamber. NativeMark protein standards were diluted 1:8 in filtered buffer (20 mM Tris pH 8.0, 100 mM NaCl, 5% glycerol, 2 mM DTT) and used to calibrate the instrument, with data acquired in droplet dilution mode and analyzed in DiscoverMP to confirm expected contrast distributions. For sample measurements, PABPC1, CSDE1, and oligo(A_30_) were prepared at 1 µM, 2 µM, and 1 µM, respectively, in filtered buffer, and 2 µL of each sample was added to 18 µL of buffer on the coverslip and mixed by pipetting immediately prior to acquisition. Measurements were collected in AcquireMP software with a 60 sec movie recorded immediately after the dilution and analyzed in DiscoverMP using the same pipeline as standards, with calibration curves applied to determine molecular masses.

### In vitro deadenylation

Deadenylation reactions were carried out in a buffer containing 20 mM PIPES/NaOH, pH 7.0, 40 mM NaCl, 10 mM KCl, and 2 mM Mg(OAc)_2_ at 37 °C. To the 5′-fluorescein-labeled UCUAAAU-A_80_ RNA (biomers.net, Ulm, Germany) substrate (50 nM) were added PABPC1 (150 nM) and CSDE1 variants (7.5 μM), and incubated at room temperature for 15 minutes. To start the reaction, 25 nM of reconstituted eight-subunit human CCR4–NOT was added. To stop the reaction at the corresponding time point, three times the reaction volume of RNA loading dye was added (95% [v/v] deionized formamide, 17.5 mM EDTA, pH 8.0, 0.01% [w/v] bromophenol blue). The products were resolved on a denaturing TBE-urea polyacrylamide gel, which was subsequently imaged using Sapphire FL Biomolecular Imager (Azure Biosystems).

### PAL-seq

Sequencing of mRNA poly(A) tail lengths in mock and MHV-infected cells was performed with PAL-seq v4 as described previously^11^. For each sample, poly(A)-tail length spike-in standards as well as poly(A)-selected mRNA from the zebrafish ZF4 cell line (0.1 ng per µg total RNA) were added to RNA samples to enable additional assessment of tail-length measurement reproducibility and normalization of tail-length abundances across samples. The poly(A)-tail length was determined using a Hidden Markov Model trained on 1% of the filtered read clusters (but no more than 50,000 and no less than 5000) randomly picked for each library as described previously^96^. For comparison of poly(A)-tail length distributions, the contribution of non-infected cells (determined by IF of nsp9 in parallel samples) was corrected by subtracting one-third of the mock-infected distribution from MHV-A59 samples or one-quarter of the mock-infected distribution from MHV-Δ99 samples. To quantify average changes in tail-length abundance per gene, we analyzed genes with a minimum of 100 poly(A) reads (or 50 reads for measurements in cells expressing nsp1). Counts were binned into 10-nt bins according to tail length, and the changes for each bin were compared across experimental conditions.

### RNA-seq

Three synthetic luciferase mRNAs were spiked into each sample for use in normalization across experimental conditions. RNA-seq libraries were generated from rRNA-depleted (NEB) total RNA using the NEXTflex Rapid Directional mRNA-seq Kit (Bioo Scientific, 5138–10). Sequencing was carried out on an Illumina HiSeq 2500 platform with 40 cycles. Reads were aligned to the reference genome using STAR (v2.4.2a) with the following options: --runMode alignReads -- outFilterMultimapNmax 1 --outReadsUnmapped Fastx --outFilterType BySJout --outSAMattributes All --outSAMtype BAM SortedByCoordinate. Gene-level exon counts were obtained with htseq-count (v0.11.0).

---

**Figure S1.**
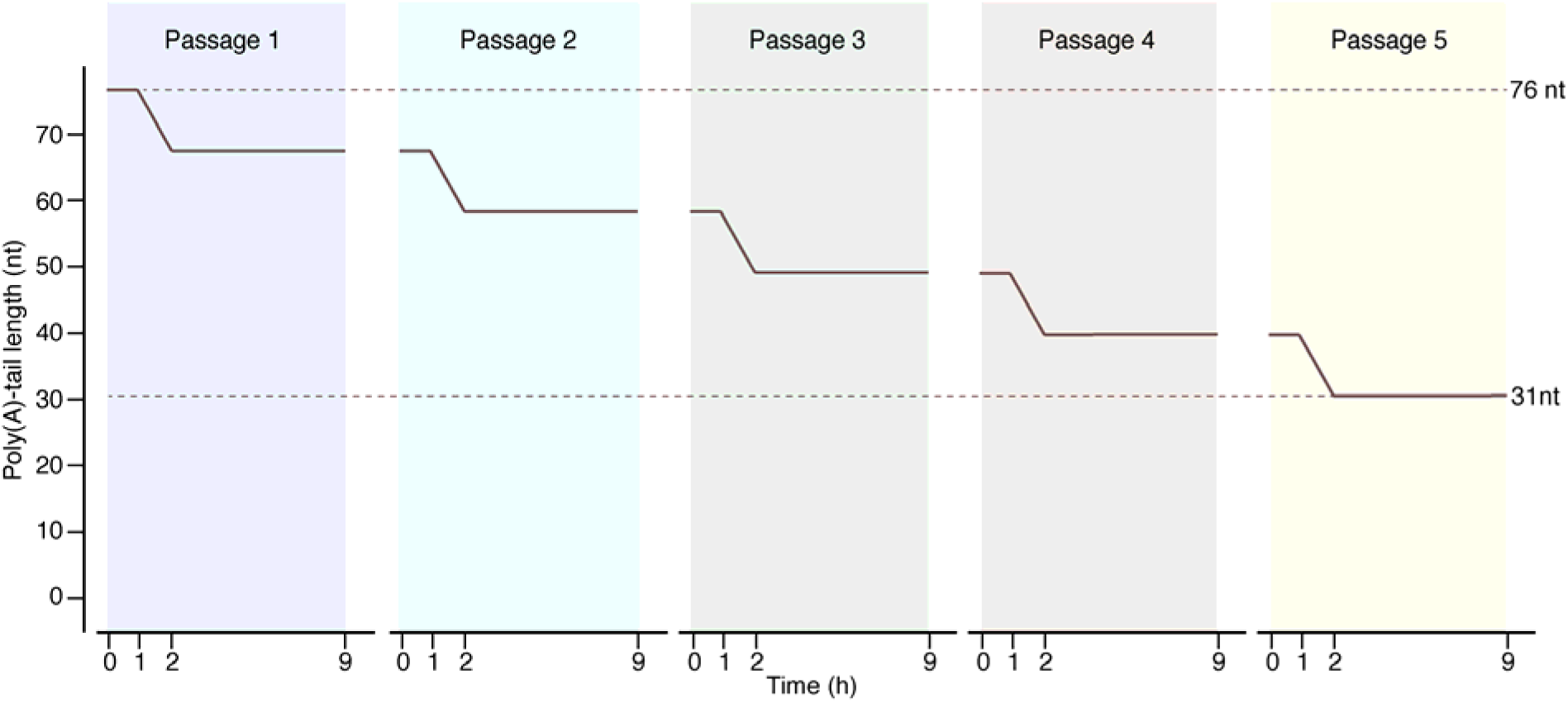
Poly(A)-tail length of viral mRNAs is expected to decrease during successive infections. A best-case estimate of poly(A)-tail length dynamics in viral mRNAs. Plotted are predicted poly(A)-tail length changes of viral mRNAs under optimal settings. During the first hour post-infection, viral mRNAs enter the cytoplasm, and their poly(A) tails remain protected within the capsid. In the second hour, as the viral genomic mRNA undergoes translation, the poly(A) tail becomes susceptible to deadenylation. In the most favorable scenario, assuming one of the slowest deadenylation rates reported for TOP mRNAs^4^ (0.15 nt/min), the tail would shorten by ∼9 nt. In the subsequent hours, once the viral RdRp is synthesized, the poly(A) tail is effectively stabilized as a poly(U) tract, and efficient encapsidation of the genomic mRNA into new virions prevents further deadenylation. Under this scheme, a genomic mRNA that begins with a 76-nt poly(A)-tail length is projected to have a 31-nt poly(A)-tail length by passage 5.

**Figure S2.**
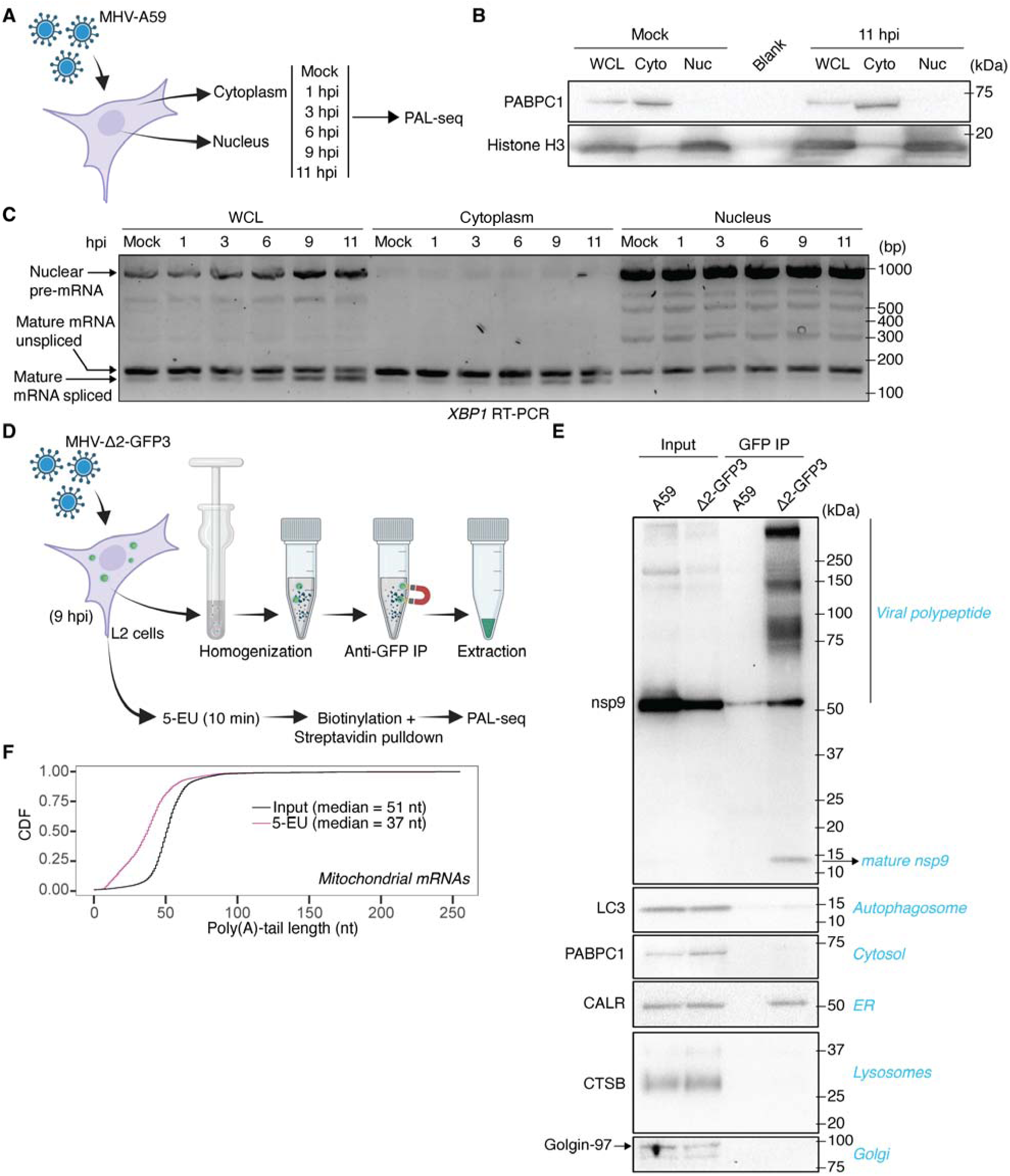
Poly(A)-tail extension of viral mRNAs occurs in double membrane vesicles. (A) Outline of infection time course and fractionation. (B) Assessment of subcellular fractionation. Shown are immunoblots of whole-cell lysates (WCL), cytoplasmic lysates (Cyto), and nuclear lysates (Nuc) of mock and infected cells (11 hpi) probed for a cytoplasmic marker (PABPC1) as well as a nuclear marker (Histone H3). (C) Assessment of subcellular fractionation by processing of Xbp1 mRNA. Shown is an agarose gel of RT-PCR products for Xbp1 mRNA in WCLs, cytoplasmic lysates, and nuclear lysates of mock or MHV-infected cells. (D) Outline of the rapid capture of DMVs adopted from organelle IP^27–29^ (top), as well as metabolic labeling for capturing the nascent mRNAs (bottom). (E) Assessment of DMV IP. Shown are immunoblots of cytoplasmic inputs and GFP IPs of cells infected with MHV-A59 or MHV-Δ2-GFP3 probed for viral protein nsp9 and various cytoplasmic markers. (F) Tail-length distributions of bulk and nascent mitochondrial mRNAs. Plotted are CDFs of poly(A)-tail lengths of total cytoplasmic and nascent mitochondrial mRNAs (input and 5-EU, respectively).

**Figure S3.**
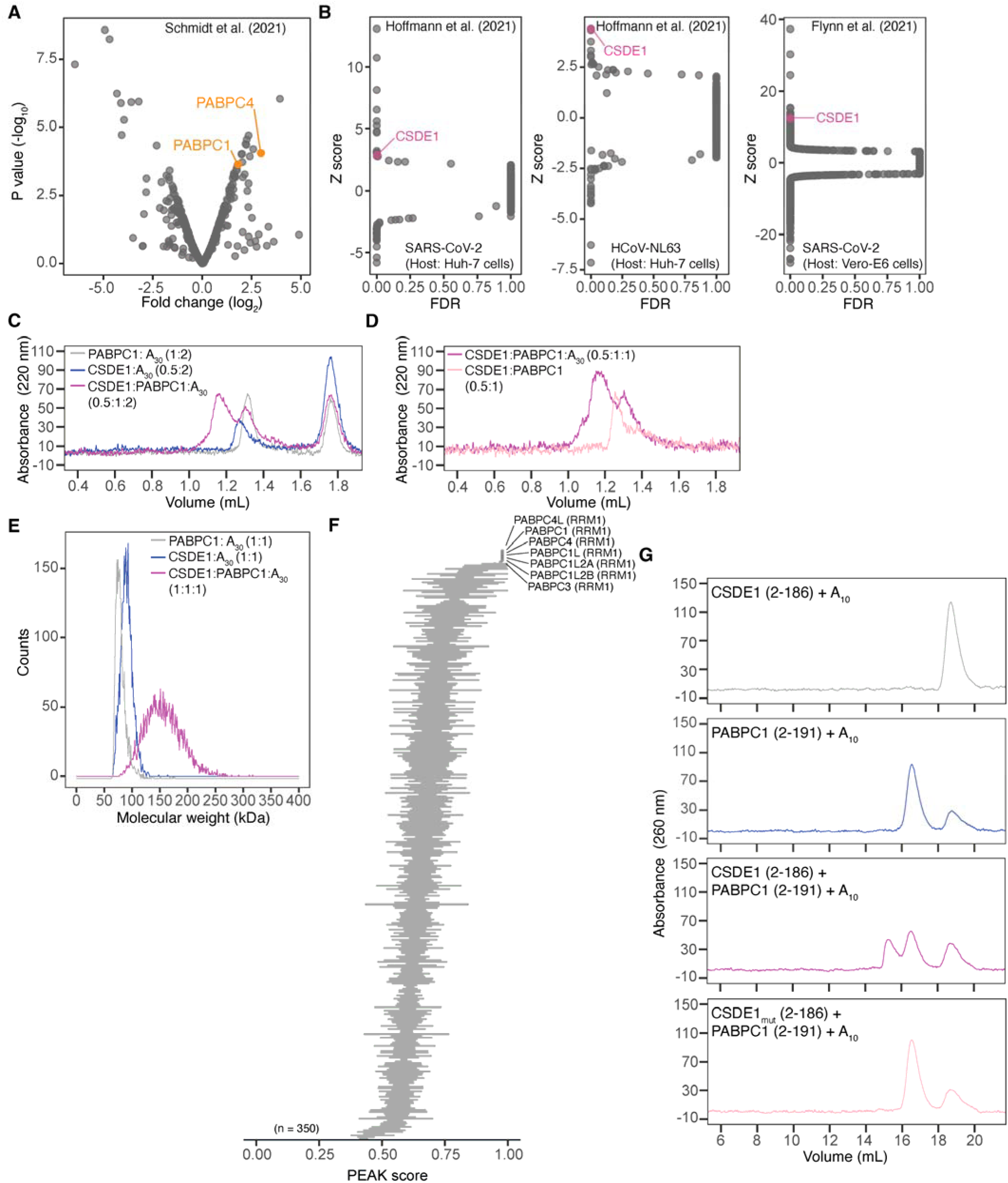
A host RBP forms a complex with viral poly(A) tail and PABPC1. (A) Replot of RAP-MS data for SARS-CoV-2 mRNAs^36^. Plotted are changes in abundance of RBPs pulled down with probes targeting SARS-CoV-2 genomic mRNA compared to proteins pulled down with probes targeting RNA component of mitochondrial RNA processing endoribonuclease (RMRP). (B) Identification of CSDE1 as a pro-viral factor. Plotted are the ranked score of CSDE1 and other proteins found in CRISPR screens designed to identify host dependencies of SARS-CoV-2 infection^48,49^. (C) Formation of a ternary complex involving full-length CSDE1, PABPC1, and the poly(A) tail. Shown are size-exclusion chromatograms of the full-length components of the CSDE1–PABPC1–oligo(A)_30_ complex. (D) Preferential binding of CSDE1 to poly(A)-bound PABPC1. Shown are size-exclusion chromatograms of CSDE1 and PABPC1 in the absence or presence of the oligo(A)_30_. (E) Stoichiometry of the ternary complex involving CSDE1, PABPC1, and oligo(A)_30_. Plotted are mass photometry profiles of the full-length components of the CSDE1–PABPC1–oligo(A)_30_ complex. (F) Predicted binding of CSD2 to the RRM1 domain of PABPC paralogs. Plotted are PEAK scores for a ColabFold^50^ screen of multimers of CSD2 of CSDE1 and all RRM domains in the human proteome. (G) Biochemical support for the AF3 model of the complex between CSDE1 and RRM1-RRM2 domain of PABPC1. Shown are size-exclusion chromatograms for absorbance at 260 nm, which enables detection of the RNA, for the experiment shown in Figure 3G.

**Figure S4.**
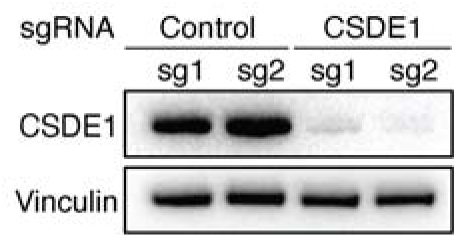
Efficiency of CSDE1 knockdown. Shown is an immunoblot probed for CSDE1 and Vinculin in control and CSDE1 KD NIH-3T3 cells targeted by one of two independent guide RNAs (sg1 and sg2).

**Figure S5.**
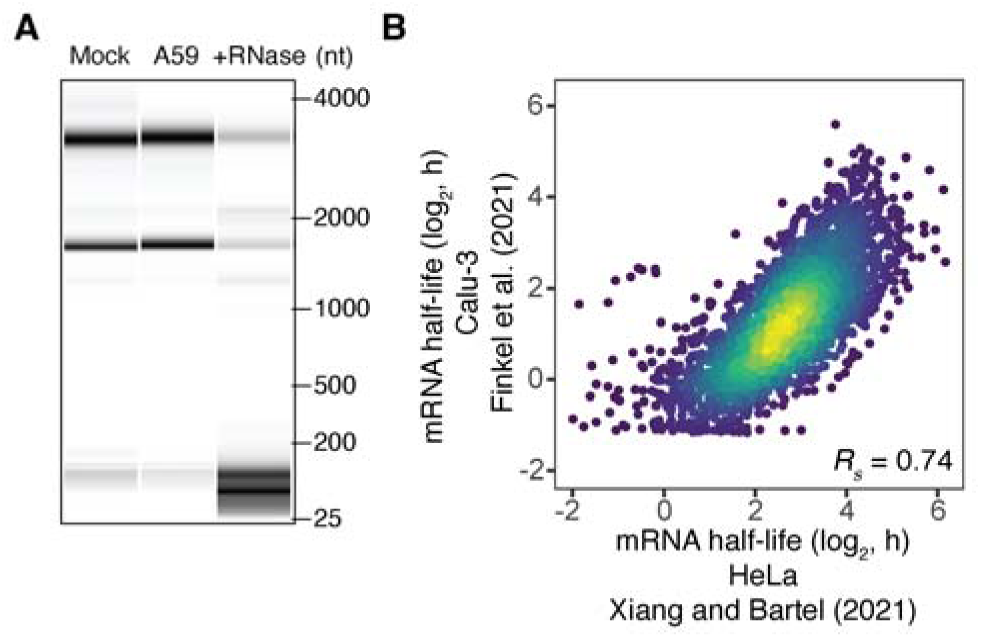
PABPC1 activity is limiting during infection, which destabilizes short-tailed host mRNAs. (A) Lack of evidence for activation of RNase L during infection with MHV-A59. Shown are bioanalyzer traces of total RNA collected from mock and MHV-infected L2 cell lysates (11 hpi) as well as from RNAse I treated lysates. (B) The correlation between half-lives of mRNAs in uninfected Calu-3 cells^57^ and in HeLa cells transfected with control siRNAs^11^.

**Figure S6.**
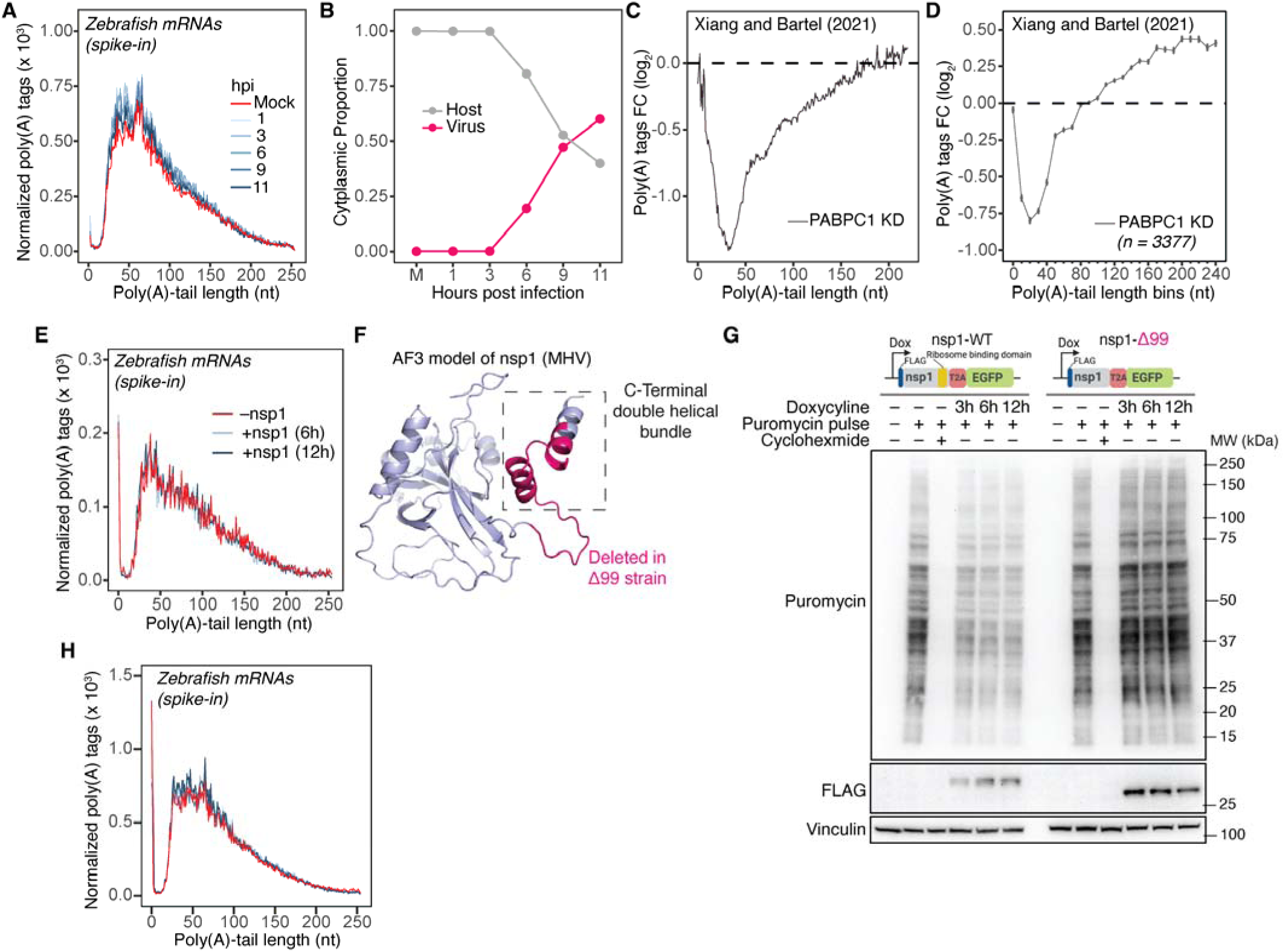
Host mRNAs with short poly(A) tails are preferentially degraded during infection. (A) Tail-length distributions of zebrafish mRNA poly(A) tags, used as spike-in controls for the experiment shown in Figure 6A. (B) Domination of host cytoplasm by viral mRNAs. Plotted are the proportion of host and viral mRNAs in the total cytoplasmic mRNA population over the course of infection. (C) Preferential degradation of short-tailed mRNAs upon PABPC1 knockdown in NIH-3T3 cells^11^. Otherwise, this panel is as in Figure 6B. (D) Preferential degradation of short-tailed mRNAs upon PABPC1 knockdown in NIH-3T3 cells^11^, analyzed after grouping mRNAs by gene; otherwise, as in Figure 6C. (E) Tail-length distributions of zebrafish mRNA poly(A) tags, used as spike-in controls for the experiment shown in Figure 6D. (F) AF3 predicted model of nsp1 from MHV. The putative ribosome-binding domain at the C terminus is indicated in a dashed box. The regions deleted in the MHV-Δ99 strain are shown in red. (G) Impact of nsp1-Δ99 on global translation levels compared to nsp1; otherwise, as in Figure 5B. (H) Tail-length distributions of zebrafish mRNA poly(A) tags, used as spike-in controls, for the experiment shown in Figure 6G.

## References

1. Edmonds, M., and Abrams, R. (1960). Polynucleotide biosynthesis: formation of a sequence of adenylate units from adenosine triphosphate by an enzyme from thymus nuclei. J. Biol. Chem. 235, 1142–1149.

2. McLaughlin, C.S., Warner, J.R., Edmonds, M., Nakazato, H., and Vaughan, M.H. (1973). Polyadenylic acid sequences in yeast messenger ribonucleic acid. J. Biol. Chem. 248, 1466– 1471.

3. Passmore, L.A., and Coller, J. (2022). Roles of mRNA poly(A) tails in regulation of eukaryotic gene expression. Nat. Rev. Mol. Cell Biol. 23, 93–106. 10.1038/s41580-021-00417-y.

4. Eisen, T.J., Eichhorn, S.W., Subtelny, A.O., Lin, K.S., McGeary, S.E., Gupta, S., and Bartel, D.P. (2020). The dynamics of cytoplasmic mRNA metabolism. Mol. Cell 77, 786–799.e10. 10.1016/j.molcel.2019.12.005.

5. Barr, J.N., and Fearns, R. (2010). How RNA viruses maintain their genome integrity. J. Gen. Virol. 91, 1373–1387. 10.1099/vir.0.020818-0.

6. van Ooij, M.J.M., Polacek, C., Glaudemans, D.H.R.F., Kuijpers, J., van Kuppeveld, F.J.M., Andino, R., Agol, V.I., and Melchers, W.J.G. (2006). Polyadenylation of genomic RNA and initiation of antigenomic RNA in a positive-strand RNA virus are controlled by the same cis-element. Nucleic Acids Res. 34, 2953–2965. 10.1093/nar/gkl349.

7. Eggen, R., Verver, J., Wellink, J., De Jong, A., Goldbach, R., and van Kammen, A. (1989). Improvements of the infectivity of in vitro transcripts from cloned cowpea mosaic virus cDNA: impact of terminal nucleotide sequences. Virology 173, 447–455. 10.1016/0042-6822(89)90557-6.

8. Neufeld, K.L., Galarza, J.M., Richards, O.C., Summers, D.F., and Ehrenfeld, E. (1994). Identification of terminal adenylyl transferase activity of the poliovirus polymerase 3Dpol. J. Virol. 68, 5811–5818. 10.1128/JVI.68.9.5811-5818.1994.

9. Tomar, S., Hardy, R.W., Smith, J.L., and Kuhn, R.J. (2006). Catalytic core of alphavirus nonstructural protein nsP4 possesses terminal adenylyltransferase activity. J. Virol. 80, 9962– 9969. 10.1128/JVI.01067-06.

10. Subtelny, A.O., Eichhorn, S.W., Chen, G.R., Sive, H., and Bartel, D.P. (2014). Poly(A)-tail profiling reveals an embryonic switch in translational control. Nature 508, 66–71. 10.1038/nature13007.

11. Xiang, K., and Bartel, D.P. (2021). The molecular basis of coupling between poly(A)-tail length and translational efficiency. eLife 10, e66493. 10.7554/eLife.66493.

12. Spector, D.H., Villa-Komaroff, L., and Baltimore, D. (1975). Studies on the function of polyadenylic acid on poliovirus RNA. Cell 6, 41–44. 10.1016/0092-8674(75)90071-9.

13. Sarnow, P. (1989). Role of 3′-end sequences in infectivity of poliovirus transcripts made in vitro. J. Virol. 63, 467–470. 10.1128/JVI.63.1.467-470.1989.

14. Silvestri, L.S., Parilla, J.M., Morasco, B.J., Ogram, S.A., and Flanegan, J.B. (2006). Relationship between poliovirus negative-strand RNA synthesis and the length of the 3′ poly(A) tail. Virology 345, 509–519. 10.1016/j.virol.2005.10.019.

15. Spagnolo, J.F., and Hogue, B.G. (2001). Requirement of the poly(A) tail in coronavirus genome replication. In The Nidoviruses: Coronaviruses and Arteriviruses, E. Lavi, S.R. Weiss, and S.T. Hingley, eds. (Springer US), 467–474. 10.1007/978-1-4615-1325-4_68.

16. Spagnolo, J.F., and Hogue, B.G. (2000). Host protein interactions with the 3′ end of bovine coronavirus RNA and the requirement of the poly(A) tail for coronavirus defective genome replication. J. Virol. 74, 5053–5065. 10.1128/JVI.74.11.5053-5065.2000.

17. Gupta, A., Li, Y., Chen, S.-H., Papas, B.N., Martin, N.P., and Morgan, M. (2023). TUT4/7-mediated uridylation of a coronavirus subgenomic RNAs delays viral replication. Commun. Biol. 6, 1040. 10.1038/s42003-023-04814-1.

18. van der Toorn, W., Bohn, P., Liu-Wei, W., Olguin-Nava, M., Gribling-Burrer, A.-S., Smyth, R.P., and von Kleist, M. (2025). Demultiplexing and barcode-specific adaptive sampling for nanopore direct RNA sequencing. Nat. Commun. 16, 3742. 10.1038/s41467-025-59102-9.

19. Kim, D., Lee, J.-Y., Yang, J.-S., Kim, J.W., Kim, V.N., and Chang, H. (2020). The architecture of SARS-CoV-2 transcriptome. Cell 181, 914–921.e10. 10.1016/j.cell.2020.04.011.

20. Yoshida, H., Matsui, T., Yamamoto, A., Okada, T., and Mori, K. (2001). XBP1 mRNA is induced by ATF6 and spliced by IRE1 in response to ER stress to produce a highly active transcription factor. Cell 107, 881–891. 10.1016/S0092-8674(01)00611-0.

21. Wu, H.-Y., Ke, T.-Y., Liao, W.-Y., and Chang, N.-Y. (2013). Regulation of coronaviral poly(A) tail length during infection. PLoS One 8, e70548. 10.1371/journal.pone.0070548.

22. Shien, J.-H., Su, Y.-D., and Wu, H.-Y. (2014). Regulation of coronaviral poly(A) tail length during infection is not coronavirus species- or host cell-specific. Virus Genes 49, 383–392. 10.1007/s11262-014-1103-7.

23. V’kovski, P., Kratzel, A., Steiner, S., Stalder, H., and Thiel, V. (2021). Coronavirus biology and replication: implications for SARS-CoV-2. Nat. Rev. Microbiol. 19, 155–170. 10.1038/s41579-020-00468-6.

24. Knoops, K., Kikkert, M., van den Worm, S.H.E., Zevenhoven-Dobbe, J.C., van der Meer, Y., Koster, A.J., Mommaas, A.M., and Snijder, E.J. (2008). SARS-coronavirus replication is supported by a reticulovesicular network of modified endoplasmic reticulum. PLoS Biol. 6, e226. 10.1371/journal.pbio.0060226.

25. Gosert, R., Kanjanahaluethai, A., Egger, D., Bienz, K., and Baker, S.C. (2002). RNA replication of mouse hepatitis virus takes place at double-membrane vesicles. J. Virol. 76, 3697–3708. 10.1128/JVI.76.8.3697-3708.2002.

26. Lai, M.M., Brayton, P.R., Armen, R.C., Patton, C.D., Pugh, C., and Stohlman, S.A. (1981). Mouse hepatitis virus A59: mRNA structure and genetic localization of the sequence divergence from hepatotropic strain MHV-3. J. Virol. 39, 823–834. 10.1128/JVI.39.3.823-834.1981.

27. Abu-Remaileh, M., Wyant, G.A., Kim, C., Laqtom, N.N., Abbasi, M., Chan, S.H., Freinkman, E., and Sabatini, D.M. (2017). Lysosomal metabolomics reveals V-ATPase and mTOR-dependent regulation of amino acid efflux from lysosomes. Science 358, 807–813. 10.1126/science.aan6298.

28. Chen, W.W., Freinkman, E., Wang, T., Birsoy, K., and Sabatini, D.M. (2016). Absolute quantification of matrix metabolites reveals the dynamics of mitochondrial metabolism. Cell 166, 1324–1337.e11. 10.1016/j.cell.2016.07.040.

29. Ray, G.J., Boydston, E.A., Shortt, E., Wyant, G.A., Lourido, S., Chen, W.W., and Sabatini, D.M. (2020). A PEROXO-tag enables rapid isolation of peroxisomes from human cells. iScience 23, 101109. 10.1016/j.isci.2020.101109.

30. Freeman, M.C., Graham, R.L., Lu, X., Peek, C.T., and Denison, M.R. (2014). Coronavirus replicase–reporter fusions provide quantitative analysis of replication and replication complex formation. J. Virol. 88, 5319–5327. 10.1128/JVI.00021-14.

31. Huang, Y., Wang, T., Zhong, L., Zhang, W., Zhang, Y., Yu, X., Yuan, S., and Ni, T. (2024). Molecular architecture of coronavirus double-membrane vesicle pore complex. Nature 633, 224–231. 10.1038/s41586-024-07817-y.

32. Wolff, G., Limpens, R.W.A.L., Zevenhoven-Dobbe, J.C., Laugks, U., Zheng, S., de Jong, A.W.M., Koning, R.I., Agard, D.A., Grünewald, K., Koster, A.J., et al. (2020). A molecular pore spans the double membrane of the coronavirus replication organelle. Science 369, 1395–1398. 10.1126/science.abd3629.

33. Chen, A., Lupan, A.-M., Quek, R.T., Stanciu, S.G., Asaftei, M., Stanciu, G.A., Hardy, K.S., de Almeida Magalhães, T., Silver, P.A., Mitchison, T.J., et al. (2024). A coronaviral pore–replicase complex links RNA synthesis and export from double-membrane vesicles. Sci. Adv. 10, eadq9580. 10.1126/sciadv.adq9580.

34. McShane, E., Couvillion, M., Ietswaart, R., Prakash, G., Smalec, B.M., Soto, I., Baxter-Koenigs, A.R., Choquet, K., and Churchman, L.S. (2024). A kinetic dichotomy between mitochondrial and nuclear gene expression processes. Mol. Cell 84, 1541–1555.e11. 10.1016/j.molcel.2024.02.028.

35. Lee, S., Lee, Y.-S., Choi, Y, Son, A., Park, Y., Lee, K.-M., Kim, J., Kim, J.-S., and Kim, V.N. (2021). The SARS-CoV-2 RNA interactome. Mol. Cell 81, 2838–2850.e6. 10.1016/j.molcel.2021.04.022.

36. Schmidt, N., Lareau, C.A., Keshishian, H., Ganskih, S., Schneider, C., Hennig, T., Melanson, R., Werner, S., Wei, Y., Zimmer, M., et al. (2021). The SARS-CoV-2 RNA–protein interactome in infected human cells. Nat. Microbiol. 6, 339–353. 10.1038/s41564-020-00846-z.

37. Webster, M.W., Chen, Y.-H., Stowell, J.A.W., Alhusaini, N., Sweet, T., Graveley, B.R., Coller, J., and Passmore, L.A. (2018). mRNA deadenylation is coupled to translation rates by the differential activities of Ccr4–Not nucleases. Mol. Cell 70, 1089–1100.e8. 10.1016/j.molcel.2018.05.033.

38. Yi, H., Park, J., Ha, M., Lim, J., Chang, H., and Kim, V.N. (2018). PABP cooperates with the CCR4–NOT complex to promote mRNA deadenylation and block precocious decay. Mol. Cell 70, 1081–1088.e5. 10.1016/j.molcel.2018.05.009.

39. Abramson, J., Adler, J., Dunger, J., Evans, R., Green, T., Pritzel, A., Ronneberger, O., Willmore, L., Ballard, A.J., Bambrick, J., et al. (2024). Accurate structure prediction of biomolecular interactions with AlphaFold 3. Nature 630, 493–500. 10.1038/s41586-024-07487-w.

40. Hohmann, U., Graf, M., Schellhaas, U., Pacheco-Fiallos, B., Fin, L., Riabov-Bassat, D., Pühringer, T., Szalay, M.-F., Tirián, L., Handler, D., et al. (2024). A molecular switch orchestrates the nuclear export of human messenger RNA. Preprint at bioRxiv. 10.1101/2024.03.24.586400.

41. Mattijssen, S., Kozlov, G., Gaidamakov, S., Ranjan, A., Fonseca, B.D., Gehring, K., and Maraia, R.J. (2021). The isolated La-module of LARP1 mediates 3′ poly(A) protection and mRNA stabilization, dependent on its intrinsic PAM2 binding to PABPC1. RNA Biol. 18, 275–289. 10.1080/15476286.2020.1860376.

42. Jiménez-López, D., and Guzmán, P. (2014). Insights into the evolution and domain structure of ataxin-2 proteins across eukaryotes. BMC Res. Notes 7, 453. 10.1186/1756-0500-7-453.

43. Boeynaems, S., Dorone, Y., Marian, A., Shabardina, V., Huang, G., Kim, G., Sanyal, A., Şen, N.-E., Docampo, R., Ruiz-Trillo, I., et al. (2021). Poly(A)-binding protein is an ataxin-2 chaperone that emulsifies biomolecular condensates. Preprint at bioRxiv. 10.1101/2021.08.23.457426.

44. Key, J., Almaguer-Mederos, L.-E., Kandi, A.R., Sen, N.-E., Gispert, S., Köpf, G., Meierhofer, D., and Auburger, G. (2025). ATXN2L primarily interacts with NUFIP2; the absence of ATXN2L results in NUFIP2 depletion, and the ATXN2 polyQ expansion triggers NUFIP2 accumulation. Neurobiol. Dis. 209, 106903. 10.1016/j.nbd.2025.106903.

45. Kozlov, G., Ménade, M., Rosenauer, A., Nguyen, L., and Gehring, K. (2010). Molecular determinants of PAM2 recognition by the MLLE domain of poly(A)-binding protein. J. Mol. Biol. 397, 397–407. 10.1016/j.jmb.2010.01.032.

46. Xie, J., Kozlov, G., and Gehring, K. (2014). The “tale” of poly(A)-binding protein: the MLLE domain and PAM2-containing proteins. Biochim. Biophys. Acta Gene Regul. Mech. 1839, 1062–1068. 10.1016/j.bbagrm.2014.08.001.

47. Jiang, L., Xiao, M., Liao, Q.-Q., Zheng, L., Li, C., Liu, Y., Yang, B., Ren, A., Jiang, C., and Feng, X.-H. (2023). High-sensitivity profiling of SARS-CoV-2 noncoding region–host protein interactome reveals the potential regulatory role of negative-sense viral RNA. mSystems 8, e00135–23. 10.1128/msystems.00135-23.

48. Hoffmann, H.-H., Sánchez-Rivera, F.J., Schneider, W.M., Luna, J.M., Soto-Feliciano, Y.M., Ashbrook, A.W., Pen, J.L., Leal, A.A., Ricardo-Lax, I., Michailidis, E., et al. (2021). Functional interrogation of a SARS-CoV-2 host protein interactome identifies unique and shared coronavirus host factors. Cell Host Microbe 29, 267–280.e5. 10.1016/j.chom.2020.12.009.

49. Flynn, R.A., Belk, J.A., Qi, Y., Yasumoto, Y., Wei, J., Alfajaro, M.M., Shi, Q., Mumbach, M.R., Limaye, A., DeWeirdt, P.C., et al. (2021). Discovery and functional interrogation of SARS-CoV-2 RNA–host protein interactions. Cell 184, 2394–2411.e16. 10.1016/j.cell.2021.03.012.

50. Kim, G., Lee, S., Levy Karin, E., Kim, H., Moriwaki, Y., Ovchinnikov, S., Steinegger, M., and Mirdita, M. (2025). Easy and accurate protein structure prediction using ColabFold. Nat. Protoc. 20, 620–642. 10.1038/s41596-024-01060-5.

51. Grosset, C., Chen, C.-Y.A., Xu, N., Sonenberg, N., Jacquemin-Sablon, H., and Shyu, A.-B. (2000). A mechanism for translationally coupled mRNA turnover: interaction between the poly(A) tail and a c-fos RNA coding determinant via a protein complex. Cell 103, 29–40. 10.1016/S0092-8674(00)00102-1.

52. Chang, T.-C., Yamashita, A., Chen, C.-Y.A., Yamashita, Y., Zhu, W., Durdan, S., Kahvejian, A., Sonenberg, N., and Shyu, A.-B. (2004). UNR, a new partner of poly(A)-binding protein, plays a key role in translationally coupled mRNA turnover mediated by the c-fos major coding-region determinant. Genes Dev. 18, 2010–2023. 10.1101/gad.1219104.

53. Schäfer, I.B., Yamashita, M., Schuller, J.M., Schüssler, S., Reichelt, P., Strauss, M., and Conti, E. (2019). Molecular basis for poly(A) RNP architecture and recognition by the Pan2–Pan3 deadenylase. Cell 177, 1619–1631.e21. 10.1016/j.cell.2019.04.013.

54. Fisher, T., Gluck, A., Narayanan, K., Kuroda, M., Nachshon, A., Hsu, J.C., Halfmann, P.J., Yahalom-Ronen, Y., Tamir, H., Finkel, Y., et al. (2022). Parsing the role of NSP1 in SARS-CoV-2 infection. Cell Rep. 40, 110954. 10.1016/j.celrep.2022.110954.

55. Shehata, S.I., and Parker, R. (2023). SARS-CoV-2 Nsp1-mediated mRNA degradation requires mRNA interaction with the ribosome. RNA Biol. 20, 444–456. 10.1080/15476286.2023.2231280.

56. Mendez, A.S., Ly, M, González-Sánchez, A.M., Hartenian, E., Ingolia, N.T., Cate, J.H., and Glaunsinger, B.A. (2021). The N-terminal domain of SARS-CoV-2 nsp1 plays key roles in suppression of cellular gene expression and preservation of viral gene expression. Cell Rep. 37, 109841. 10.1016/j.celrep.2021.109841.

57. Finkel, Y., Gluck, A., Nachshon, A., Winkler, R., Fisher, T., Rozman, B., Mizrahi, O., Lubelsky, Y., Zuckerman, B., Slobodin, B., et al. (2021). SARS-CoV-2 uses a multipronged strategy to impede host protein synthesis. Nature 594, 240–245. 10.1038/s41586-021-03610-3.

58. Schubert, K., Karousis, E.D., Ban, I., Lapointe, C., Leibundgut, M., Baeumlin, E., Kummerant, E., Scaiola, A., Schoenhut, T., Ziegelmueller, J., et al. (2023). Universal features of Nsp1-mediated translational shutdown by coronaviruses. Preprint at bioRxiv. 10.1101/2023.05.31.543022.

59. Schubert, K., Karousis, E.D., Jomaa, A., Scaiola, A., Echeverria, B., Gurzeler, L.-A., Leibundgut, M., Thiel, V., Mühlemann, O., and Ban, N. (2020). SARS-CoV-2 Nsp1 binds the ribosomal mRNA channel to inhibit translation. Nat. Struct. Mol. Biol. 27, 959–966. 10.1038/s41594-020-0511-8.

60. Baer, B.W., and Kornberg, R.D. (1980). Repeating structure of cytoplasmic poly(A)-ribonucleoprotein. Proc. Natl. Acad. Sci. USA 77, 1890–1892. 10.1073/pnas.77.4.1890.

61. Smith, B.L., Gallie, D.R., Le, H., and Hansma, P.K. (1997). Visualization of poly(A)-binding protein complex formation with poly(A) RNA using atomic force microscopy. J. Struct. Biol. 119, 109–117. 10.1006/jsbi.1997.3864.

62. Lin, J., Fabian, M., Sonenberg, N., and Meller, A. (2012). Nanopore detachment kinetics of poly(A)-binding proteins from RNA molecules reveals the critical role of C-terminus interactions. Biophys. J. 102, 1427–1434. 10.1016/j.bpj.2012.02.025.

63. Yoshida, M., Yoshida, K., Kozlov, G., Lim, N.S., De Crescenzo, G., Pang, Z., Berlanga, J.J., Kahvejian, A., Gehring, K., Wing, S.S., et al. (2006). Poly(A)-binding protein (PABP) homeostasis is mediated by the stability of its inhibitor, Paip2. EMBO J. 25, 1934–1944. 10.1038/sj.emboj.7601079.

64. Williams, R.K., Jiang, G.S., and Holmes, K.V. (1991). Receptor for mouse hepatitis virus is a member of the carcinoembryonic antigen family of glycoproteins. Proc. Natl. Acad. Sci. USA 88, 5533–5536. 10.1073/pnas.88.13.5533.

65. Dveksler, G.S., Pensiero, M.N., Cardellichio, C.B., Williams, R.K., Jiang, G.S., Holmes, K.V., and Dieffenbach, C.W. (1991). Cloning of the mouse hepatitis virus (MHV) receptor: expression in human and hamster cell lines confers susceptibility to MHV. J. Virol. 65, 6881–6891. 10.1128/JVI.65.12.6881-6891.1991.

66. Iwamoto, M., Björklund, T., Lundberg, C., Kirik, D., and Wandless, T.J. (2010). A general chemical method to regulate protein stability in the mammalian central nervous system. Chem. Biol. 17, 981–988. 10.1016/j.chembiol.2010.07.009.

67. Züst, R., Cervantes-Barragán, L., Kuri, T., Blakqori, G., Weber, F., Ludewig, B., and Thiel, V. (2007). Coronavirus non-structural protein 1 is a major pathogenicity factor: implications for the rational design of coronavirus vaccines. PLoS Pathog. 3, e109. 10.1371/journal.ppat.0030109.

68. Brinton, M.A., and Basu, M. (2015). Functions of the 3′ and 5′ genome RNA regions of members of the genus Flavivirus. Virus Res. 206, 108–119. 10.1016/j.virusres.2015.02.006.

69. Brouze, M., Czarnocka-Cieciura, A., Gewartowska, O., Kusio-Kobiałka, M., Jachacy, K., Szpila, M., Tarkowski, B., Gruchota, J., Krawczyk, P., Mroczek, S., et al. (2024). TENT5-mediated polyadenylation of mRNAs encoding secreted proteins is essential for gametogenesis in mice. Nat. Commun. 15, 5331. 10.1038/s41467-024-49479-4.

70. Kim, D., Lee, Y., Jung, S.-J., Yeo, J., Seo, J.J., Lee, Y.-Y., Lim, J., Chang, H., Song, J., Yang, J., et al. (2020). Viral hijacking of the TENT4–ZCCHC14 complex protects viral RNAs via mixed tailing. Nat. Struct. Mol. Biol. 27, 581–588. 10.1038/s41594-020-0427-3.

71. Lee, Y.-S., Levdansky, Y., Jung, Y., Kim, V.N., and Valkov, E. (2024). Deadenylation kinetics of mixed poly(A) tails at single-nucleotide resolution. Nat. Struct. Mol. Biol. 31, 826–834. 10.1038/s41594-023-01187-1.

72. Lim, J., Kim, D., Lee, Y., Ha, M., Lee, M., Yeo, J., Chang, H., Song, J., Ahn, K., and Kim, V.N. (2018). Mixed tailing by TENT4A and TENT4B shields mRNA from rapid deadenylation. Science 361, 701–704. 10.1126/science.aam5794.

73. Seo, J.J., Jung, S.-J., Yang, J., Choi, D.-E., and Kim, V.N. (2023). Functional viromic screens uncover regulatory RNA elements. Cell 186, 3291–3306.e21. 10.1016/j.cell.2023.06.007.

74. Eisen, T.J., Li, J.J., and Bartel, D.P. (2022). The interplay between translational efficiency, poly(A) tails, microRNAs, and neuronal activation. RNA 28, 808–831. 10.1261/rna.079046.121.

75. Du, L., and Richter, J.D. (2005). Activity-dependent polyadenylation in neurons. RNA 11, 1340– 1347. 10.1261/rna.2870505.

76. Wu, L., Wells, D., Tay, J., Mendis, D., Abbott, M.-A., Barnitt, A., Quinlan, E., Heynen, A., Fallon, J.R., and Richter, J.D. (1998). CPEB-mediated cytoplasmic polyadenylation and the regulation of experience-dependent translation of α-CaMKII mRNA at synapses. Neuron 21, 1129–1139. 10.1016/S0896-6273(00)80630-3.

77. Safaee, N., Kozlov, G., Noronha, A.M., Xie, J., Wilds, C.J., and Gehring, K. (2012). Interdomain allostery promotes assembly of the poly(A) mRNA complex with PABP and eIF4G. Mol. Cell 48, 375–386. 10.1016/j.molcel.2012.09.001.

78. Kajjo, S., Sharma, S., Chen, S., Brothers, W.R., Cott, M., Hasaj, B., Jovanovic, P., Larsson, O., and Fabian, M.R. (2022). PABP prevents the untimely decay of select mRNA populations in human cells. EMBO J. 41, e108650. 10.15252/embj.2021108650.

79. Otero, L.J., Ashe, M.P., and Sachs, A.B. (1999). The yeast poly(A)-binding protein Pab1p stimulates in vitro poly(A)-dependent and cap-dependent translation by distinct mechanisms. EMBO J. 18, 3153–3163. 10.1093/emboj/18.11.3153

80. Chang, J.J.-Y., Gleeson, J., Rawlinson, D., De Paoli-Iseppi, R., Zhou, C., Mordant, F.L., Londrigan, S.L., Clark, M.B., Subbarao, K., Stinear, T.P., et al. (2022). Long-read RNA sequencing identifies polyadenylation elongation and differential transcript usage of host transcripts during SARS-CoV-2 in vitro infection. Front. Immunol. 13, 832223. 10.3389/fimmu.2022.832223.

81. Anderson, E.C., Hunt, S.L., and Jackson, R.J. (2007). Internal initiation of translation from the human rhinovirus-2 internal ribosome entry site requires the binding of Unr to two distinct sites on the 5′ untranslated region. J. Gen. Virol. 88, 3043–3052. 10.1099/vir.0.82463-0.

82. Boussadia, O., Niepmann, M., Créancier, L., Prats, A.-C., Dautry, F., and Jacquemin-Sablon, H. (2003). Unr is required in vivo for efficient initiation of translation from the internal ribosome entry sites of both rhinovirus and poliovirus. J. Virol. 77, 3353–3359. 10.1128/JVI.77.6.3353-3359.2003.

83. Hunt, S.L., Hsuan, J.J., Totty, N., and Jackson, R.J. (1999). Unr, a cellular cytoplasmic RNA-binding protein with five cold-shock domains, is required for internal initiation of translation of human rhinovirus RNA. Genes Dev. 13, 437–448. 10.1101/gad.13.4.437.

84. Aviner, R., Li, K.H., Frydman, J., and Andino, R. (2021). Cotranslational prolyl hydroxylation is essential for flavivirus biogenesis. Nature 596, 558–564. 10.1038/s41586-021-03851-2.

85. Herold, J., and Andino, R. (2001). Poliovirus RNA replication requires genome circularization through a protein–protein bridge. Mol. Cell 7, 581–591. 10.1016/S1097-2765(01)00205-2.

86. Ghanim, G.E., Hu, H., Boulanger, J., and Nguyen, T.H.D. (2025). Structural mechanism of LINE-1 target-primed reverse transcription. Science 388, eads8412. 10.1126/science.ads8412.

87. Dewannieux, M., and Heidmann, T. (2005). Role of poly(A) tail length in Alu retrotransposition. Genomics 86, 378–381. 10.1016/j.ygeno.2005.05.009.

88. Doucet, A.J., Wilusz, J.E., Miyoshi, T., Liu, Y., and Moran, J.V. (2015). A 3′ poly(A) tract is required for LINE-1 retrotransposition. Mol. Cell 60, 728–741. 10.1016/j.molcel.2015.10.012.

89. Thawani, A., Florez Ariza, A.J., Nogales, E., and Collins, K. (2024). Template and target-site recognition by human LINE-1 in retrotransposition. Nature 626, 186–193. 10.1038/s41586-023-06933-5.

90. Leibowitz, J., Kaufman, G., and Liu, P. (2011). Coronaviruses: propagation, quantification, storage, and construction of recombinant mouse hepatitis virus. Curr. Protoc. Microbiol. Chapter 15, Unit 15E.1. 10.1002/9780471729259.mc15e01s21.

91. Ray, G.J., Nardini, E., Keys, H.R., Lin, D.H., Sabatini, D.M., and Bartel, D.P. (2025). Lysosomal RNA profiling reveals targeting of specific types of RNAs for degradation. Preprint at bioRxiv. 10.1101/2025.09.09.674968.

92. Liao, J.-Y., Yang, B., Shi, C.-P., Deng, W.-X., Deng, J.-S., Cen, M.-F., Zheng, B.-Q., Zhan, Z.-L., Liang, Q.-L., Wang, J.-E., et al. (2025). RBPWorld for exploring functions and disease associations of RNA-binding proteins across species. Nucleic Acids Res. 53, D220–D232. 10.1093/nar/gkae1028.

93. Raisch, T., Chang, C.-T., Levdansky, Y., Muthukumar, S., Raunser, S., and Valkov, E. (2019). Reconstitution of recombinant human CCR4–NOT reveals molecular insights into regulated deadenylation. Nat. Commun. 10, 3173. 10.1038/s41467-019-11094-z.

94. Levdansky, Y., and Valkov, E. (2024). Reconstitution of human CCR4–NOT complex from purified proteins and an assay of its deadenylation activity. Methods Mol. Biol. 2723, 1–17. 10.1007/978-1-0716-3481-3_1.

95. Sasse, A., Ray, D., Laverty, K.U., Tam, C.L., Albu, M., Zheng, H., Levdansky, Y., Lyudovyk, O., Dalal, T., Nie, K., et al. (2025). A resource of RNA-binding protein motifs across eukaryotes reveals evolutionary dynamics and gene-regulatory function. Nat. Biotechnol., 1–11. 10.1038/s41587-025-02733-6.

96. Xiang, K., Ly, J., and Bartel, D.P. (2024). Control of poly(A)-tail length and translation in vertebrate oocytes and early embryos. Dev. Cell 59, 1058–1074.e11. 10.1016/j.devcel.2024.02.007.

